# Vitamin D signaling orchestrates skeletal muscle metabolic flexibility by regulating its fuel choice

**DOI:** 10.1101/2023.03.05.531218

**Authors:** Anamica Das, Neha Jawla, Suchitra D. Gopinath, G. Aneeshkumar Arimbasseri

## Abstract

Vitamin D deficiency is associated with skeletal muscle pathologies. However, the role of vitamin D signaling in maintenance of muscle function is not well understood. Mice lacking vitamin D receptor (VDR) exhibit severe muscle wasting after weaning and this is associated with accumulation of muscle glycogen and energy deprivation. Here we show that the skeletal muscles of vdr-/- mice exhibit upregulation of fatty acid oxidation pathway and PPAR pathway and are predisposed to utilize fatty acids as the energy source even in a carbohydrate-enriched diet. As a result, fat-enriched diets could alleviate energy deprivation and atrophy of vdr-/- skeletal muscles. However, the complete restoration of muscle mass and systemic metabolism of vdr-/- mice depended on the quality of diets. Despite increasing muscle energy levels, a lard-based high-fat diet (HFD) disrupted glucose homeostasis by specifically inhibiting the insulin synthesis in pancreatic islets. Surprisingly, milk-based high-fat diets (MBD) could restore both muscle mass and pancreatic insulin response. This study reveals a micronutrient-macronutrient interaction network that connects vitamin D signaling with muscle fuel selection and pancreatic insulin response to enable energy homeostasis under different metabolic landscapes.

## INTRODUCTION

Coordinated changes in organs and tissues such as liver, adipose, pancreas, and skeletal muscles are required for adaptation of the body to varying metabolic and nutritional landscapes to maintain homeostasis (Smith et al. 2018). As the largest insulin-sensitive tissue in the body, the inability of skeletal muscles to switch between fatty acids and glucose as energy sources during different metabolic conditions impairs metabolic flexibility and leads to disorders such as diabetes (Stump et al. 2006; Galgani, Moro, and Ravussin 2008). Chronic imbalance in the energy intake and utilization can lead to the loss of metabolic flexibility primarily by affecting skeletal muscle insulin sensitivity (Kelley and Mandarino 2000). Intake of micronutrients, especially fat-soluble vitamins, also influence the development of metabolic syndrome (da Cunha et al. 2016; Goncalves and Amiot 2017). However, the nature of the micronutrient-macronutrient interactions that maintain metabolic flexibility has yet to be understood.

Vitamin D is an essential fat-soluble vitamin and steroid hormone, which acts on several tissues, including intestinal epithelium, bones, immune cells, and the nervous system. Vitamin D binds to its cognate nuclear receptor, VDR, which regulates the transcription of genes that have a vitamin D response element (VRE) on the promoter (Kato 2000). One of vitamin D’s primary functions is to maintain calcium and phosphate homeostasis. Ligand-bound VDR activates the transcription of intestinal calcium transporters and other calcium-binding proteins essential for intestinal calcium absorption (Christakos et al. 2011). Vitamin D function is regulated at two levels. The availability of the active form of vitamin D, 1,25-dihydroxy vitamin D, is tightly regulated at the level of synthesis and degradation (Henry 2011). Moreover, VDR exhibits variable expression in different tissues, with intestinal epithelial cells, immune cells, pancreatic islets, and bones exhibiting high levels of VDR (Lee et al. 2018). However, vitamin D has a functional impact on multiple tissues irrespective of the expression levels of VDR, as many of these effects are indirect.

Severe vitamin D deficiency and lack of VDR signaling are well-established to affect skeletal muscle functions leading to hypotonia, fatigue, falls, and sarcopenia (Visser et al. 2003; Snijder et al. 2006; Shimizu et al. 2015; Janssen et al. 2013). Nevertheless, the mechanisms by which vitamin D controls muscle mass are yet to be elucidated. Multiple studies have shown the expression of VDR in skeletal muscles at different stages of development, but its levels are low, probably because of the cell-type specific expression (Girgis et al. 2014; Pike 2014). Lack of VDR or vitamin D deficiency has been shown to reduce the expression levels of genes associated with intracellular calcium mobilization, a process essential for muscle function (Vazquez and de Boland 1996; Capiati, Vazquez, and Boland 2001). Since VDR is also expressed in the nervous system, it has been proposed that the defects in neuromuscular interactions could be responsible for muscle defects observed in vitamin D deficient conditions (Minasyan et al. 2009). Vitamin D deficiency and the absence of VDR lead to the upregulation of muscle-specific E3 ubiquitin ligases, linking vitamin D deficiency with muscle atrophy (Bhat et al. 2013). Mice lacking VDR (vdr-/-) exhibit severe skeletal muscle abnormalities including, muscle fiber atrophy (Endo et al. 2003). However, myocyte or mature myofiber-specific deletions of VDR did not show severe muscle wasting observed in whole-body knockout, though they exhibited muscle weakness and sarcopenia (Girgis et al. 2015; Minasyan et al. 2009). These data indicate that vitamin D affects skeletal muscles both directly and indirectly. However, the nature of these effects are not known.

Vitamin D deficiency is associated with systemic metabolic disorders. Epidemiological data shows that hypovitaminosis D is associated with metabolic disorders such as insulin resistance, type 2 diabetes, and NAFLD (Wimalawansa 2018; Barchetta, Cimini, and Cavallo 2020). Obese individuals with vitamin D deficiency are especially susceptible to metabolic disorders, especially type 2 diabetes. Muscle is a vital metabolic organ, and its dysfunctions are associated with systemic energy metabolism dysfunction (Sartori, Romanello, and Sandri 2021). The effect of vitamin D on muscle metabolism in connection with the systemic metabolic balance is critical in understanding the muscular pathology under vitamin D deficiency. Vitamin D increases oxygen consumption and oxidative phosphorylation and decreases oxidative stress in human skeletal muscles (Ryan et al. 2016; Sinha et al. 2013; Dzik et al. 2018). In skeletal muscles of high-fat, high-sugar-fed obese mice, supplementation of vitamin D led to a decrease in body weight and improved glucose tolerance, and reduced myosteatosis (Benetti et al. 2018). Moreover, vitamin D is also known to affect adipose tissue metabolism and pancreatic insulin response, which affect muscle metabolism (Narvaez et al. 2009; Zeitz et al. 2003). In support of this, previous studies have shown that mice lacking VDR resist high-fat diet-induced obesity (Narvaez et al. 2009).

Our previous study showed that mice lacking VDR (vdr-/-) exhibit severe glycogen storage disorder in the skeletal muscles, which leads to systemic energy deprivation and severe muscle atrophy (Das, Gopinath, and Arimbasseri 2022). However, these mice do not exhibit such metabolic defects when they are in the suckling stage; milk is a fat-enriched diet (Endo et al. 2003). Skeletal muscle is a metabolically flexible tissue that can switch between glucose and fatty acid as the energy source when dietary macronutrient composition is altered (Philip J. Randle 1998; P J Randle, Newsholme, and Garland 1964). Here, we addressed if vitamin D plays any role in the metabolic flexibility of skeletal muscles when macronutrient ratios in diet are altered. Our data shows that skeletal muscles of vdr-/- mice are pre-disposed to the utilization of fatty acids as the energy source with upregulation of PDK4 and fatty acid oxidation gene expression and mitochondrial activity. Upon providing milk-based diets, which have higher fat content, we found that the energy imbalance in the vdr-/- could be reverted. The results clearly show that the ability of skeletal muscles to switch between glucose and fatty acids as the energy source is lost in the absence of vitamin D signaling. Furthermore, we also show that vitamin D signaling is essential for insulin production by pancreatic islets when the mice were subjected to a lard-based high-fat diet. Thus, the data presented here reveal complex micronutrient-macronutrient interactions that govern metabolic flexibility and homeostasis.

## RESULTS

### vdr-/- skeletal muscles exhibit upregulation of genes associated with fatty acid metabolism.

To identify how the absence of vitamin D receptor (VDR) affects various pathways and processes at the transcriptome level, we performed RNA sequencing (RNAseq) analysis of the quadriceps (QUAD) muscle of 7-week-old wild-type (WT) and vdr-/- mice. They exhibited drastic differences in the global gene expression landscape; 287 genes were upregulated, and 89 genes were downregulated in vdr-/- (p adjusted <0.1 and |log2(fold change)| >0.6; Figure 1A, Supplementary Data Set 1).

**Figure 1.**
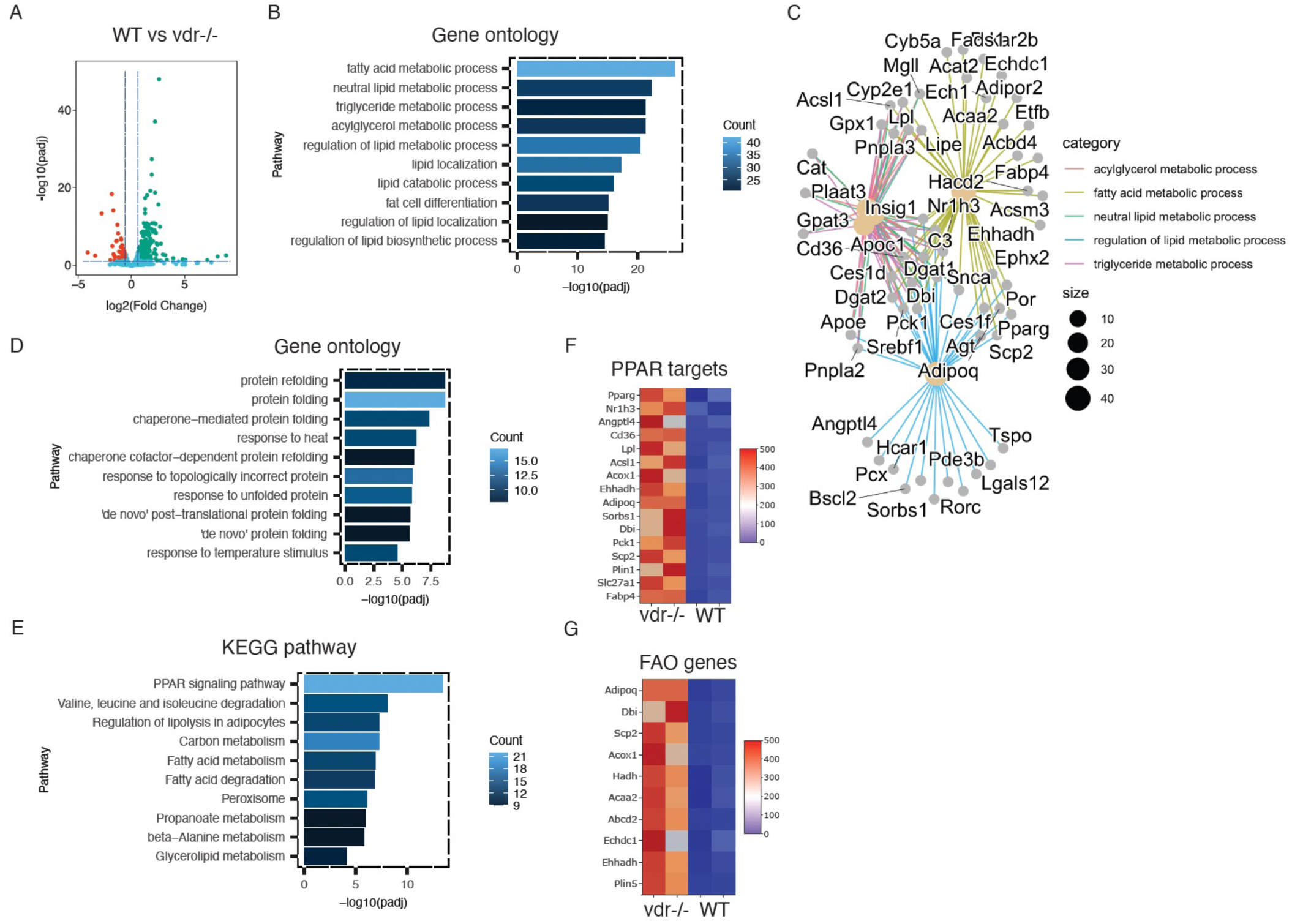
vdr-/- skeletal muscles are predisposed for fatty acid catabolism. **A**. Volcano plot highlighting the mRNAs that are differentially expressed in WT and vdr-/- QUAD muscles. 287 genes were upregulated (green) and 89 gene were downregulated (red). The blue dotted lines indicate cut-off values (p adjusted <0.1 and |log2(fold change)| & >0.6. **B, D & E**. Barplot showing -log10(padj) values of GO terms enriched among genes upregulated in vdr-/- (B), downregulated in vdr-/- (D), and KEGG pathways enriched among upregulated genes. Color scale indicate the number of term-associated genes in the upregulated gene set. **C**. A network plot showing the genes (smaller dots) associated with different lipid metabolic pathways (hubs) enriched in vdr-/- upregulated genes. Edge colors indicate the identity of pathway and the hub size indicate the number of term-associated genes. **F&G**. Heatmaps showing the normalized read counts of PPAR pathway genes (F) and fatty acid oxidation associated genes (G).

Gene ontology analysis of the genes upregulated in the vdr-/- muscle transcriptome revealed enrichment of pathways involved in fatty acid and lipid metabolism (Figure 1B&C). On the other hand, the downregulated genes show enrichment for functions associated with protein folding (Figure 1D and Supplementary Figure 1A). KEGG pathway analysis revealed that the amino acid (valine, leucine, and isoleucine) degradation pathway and multiple pathways associated with lipid and carbon metabolism are enriched among the upregulated genes (Figure 1E).

The PPAR signaling pathway is one of the critical regulators of lipid metabolism and energy metabolism. Three PPAR family members, PPAR*α*, PPAR*δ*, and PPAR*γ*, regulate different facets of lipid biogenesis and oxidation (Gervois et al. 2000). The genes upregulated in vdr-/- muscles are enriched for PPAR pathway genes (Figure 1F), primarily associated with fatty acid transport and oxidation (Supplementary Figure 1B). Interestingly, several genes directly involved in fatty acid beta-oxidation, such as Ech1, Ehhadh, Hhdh, and Acaa2, are upregulated in vdr-/- (Figure 1G and Supplementary Figure 1C). Moreover, genes associated with peroxisomal fatty acid oxidation, a process essential for energy production from long-chain fatty acids, are also upregulated in vdr-/- mice (Supplementary Figure 1D). Previously, we have shown that the absence of VDR leads to a carbohydrate utilization defect due to increased glycogen accumulation in vdr-/- muscles, causing muscle energy deficiency and an increase in their mitochondrial activity (Das, Gopinath, and Arimbasseri 2022). We hypothesized that these changes indicate the adaptation of these muscles more towards utilizing fatty acids as the energy source to circumvent defective carbohydrate metabolism.

### Milk-like diets ameliorate skeletal muscle atrophy in vdr-/- mice

When subjected to diets enriched in fat, muscles utilize fatty acids as their primary energy source, sparing glucose (P J Randle, Newsholme, and Garland 1964; P. J. Randle et al. 1963). Moreover, vdr-/- mice did not exhibit any muscle-wasting phenotype during the suckling stage when milk is the primary mode of nutrition; the fat-enriched milk diet could prevent phenotype onset in the suckling stage mice (Endo et al. 2003). To test whether diet-driven alternative pathways for muscle energy homeostasis can bypass the metabolic defects observed in vdr-/- mice, we used two diets based on milk. The first is a custom-designed nutrient formula mimicking milk composition (denoted as MFD), and the second is a commercially available milk-based formula (Nestle Lactogen 2, denoted as MBD). Both diets had a higher fat content than chow (Supplementary Table 2), and the source of fat was butter. These diets differed significantly in their fat to carbohydrate ratio (chow = 0.4, MBD = 0.76, MFD = 3). The calcium level of MFD was comparable to chow (MFD: 0.9% vs. chow: 0.7), while the calcium concentration of MBD was lower (MBD: 0.45% vs. Chow: 0.7%). Both diets have much lower calcium than the VDR rescue diet (2% calcium) used to correct mineral homeostasis in vdr-/- mice (Song, Kato, and Fleet 2003).

After weaning, WT and vdr-/- mice were allowed to feed ad libitum on MFD, MBD or chow until 7 weeks of age. Both MFD and MBD restored the growth of vdr-/- mice, as indicated by the increased body size (Figure 2A and Supplementary Figure 2A). In agreement with this, vdr-/- mice on these diets maintained body weights indistinguishable from WT mice at 5, 6, and 7 weeks of age (Figure 2B and Supplementary Figure 2B). Tibialis anterior (TA) and quadriceps (QUAD) muscle weights were restored in mice fed with MFD and MBD, indicating abrogation of muscle atrophy in vdr-/- mice (Figure 2C & D, and Supplementary Figure 2C). Histopathology analysis of the GAS and TA muscle cryo-sections revealed reduced muscle fiber cross section area in vdr-/- on chow, confirming severe atrophy, while vdr-/- on MBD diet had normal cross section area (Figure 2E). Quantifying the cross-sectional area of muscle fibers shows that vdr-/- on chow has a higher percentage of fibers with a cross-sectional area of 100-300µm2 while the majority of the fibers in WT and vdr-/- on MBD have a cross-sectional area higher than 300µm2 (Supplementary Figure 2D). These results show that despite having a difference in absolute fat content, both milk-based diets could restore muscle mass. Since the effect of both diets on growth and muscle mass were similar, for most of the further studies we used MBD with key experiments reproduced in MFD.

**Figure 2:**
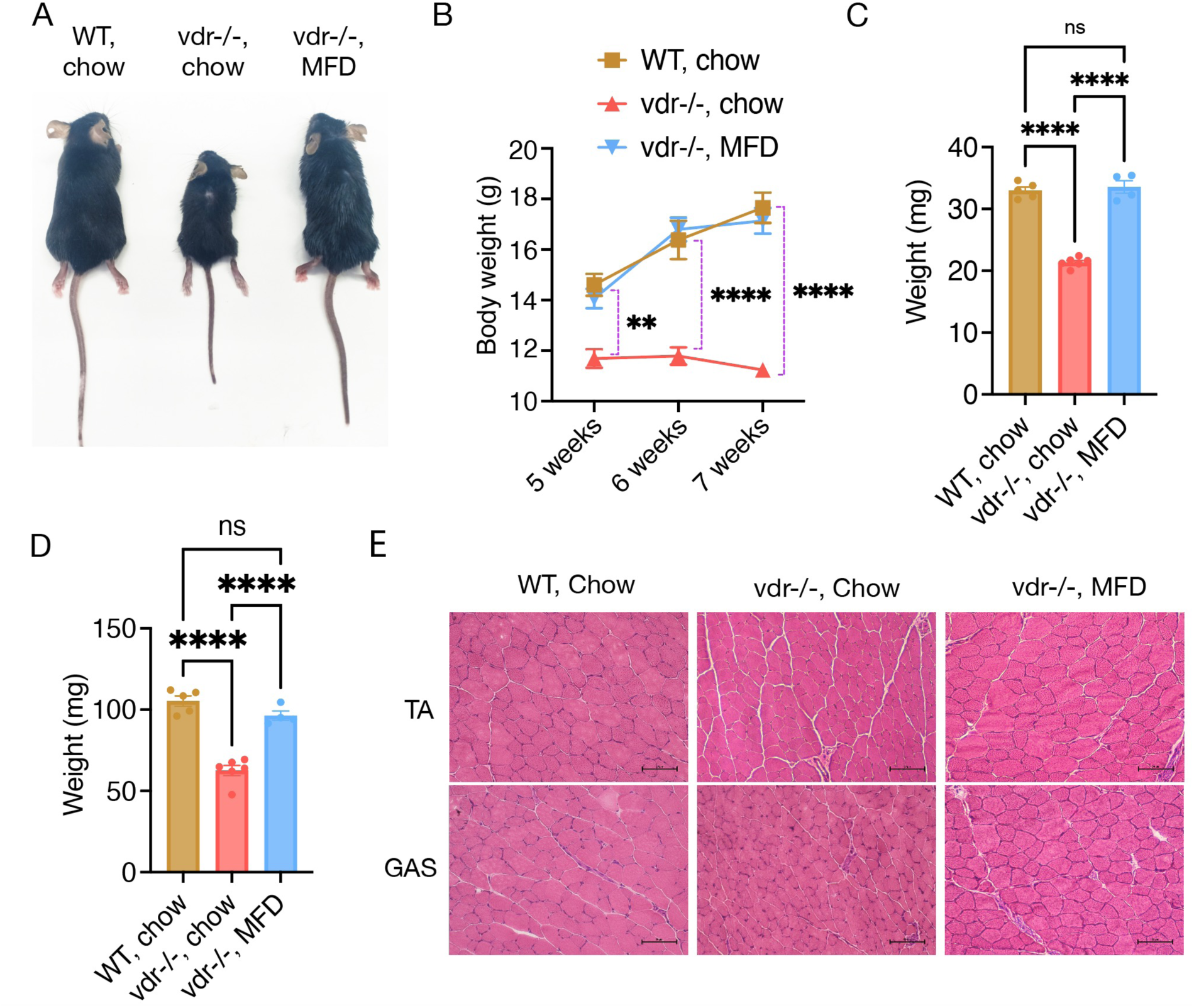
Milk based diets restore skeletal muscle atrophy in vdr-/- mice. **A.** Representative image displaying body size of WT and vdr-/- mice fed on chow, and MFD diets. **B.** Body weight (in grams) of WT and vdr−/− mice at 5, 6 and 7 weeks of age on Chow and MFD diets (*n* > 5). **C &D.** Tibialis anterior (C), and QUAD (D) muscle weight of 7-week-old WT and vdr−/− on Chow and MFD diets. **E.** Representative images of H&E stained transverse sections of gastrocnemius (GAS) muscles of WT and vdr−/− mice at 7 weeks of age on Chow and MFD diet. Graphs show mean ± SEM. *p < 0.05, **p < 0.01, ***p < 0.001, ****p < 0.0001 by two-way ANOVA (B) or by one-way ANOVA (C and D). Number of samples are denoted by the dots in the graphs.

### Milk-fat enriched diet restores the protein and energy homeostasis in skeletal muscles

The energy metabolism of muscle fibers is an important determinant of muscle mass. So, we asked if vdr-/- mice exhibit any difference in the muscle energy status. As expected, vdr-/- exhibited lower ATP levels on chow, but it was restored on MBD diet (Figure 3A), indicating that the muscle energy deficiency is alleviated on MBD. In accordance with this, the AMPK pathway, the sensor of energy deficiency in the cells, is upregulated in vdr-/- on chow, while milk-based diets reduced the levels of both LKB1 and p-AMPK, the upstream activator and effector kinases of AMPK pathway, respectively (Figure 3B & Supplementary Figure 3A). This data indicates restoration of skeletal muscle energy levels by milk-based diets.

**Figure 3:**
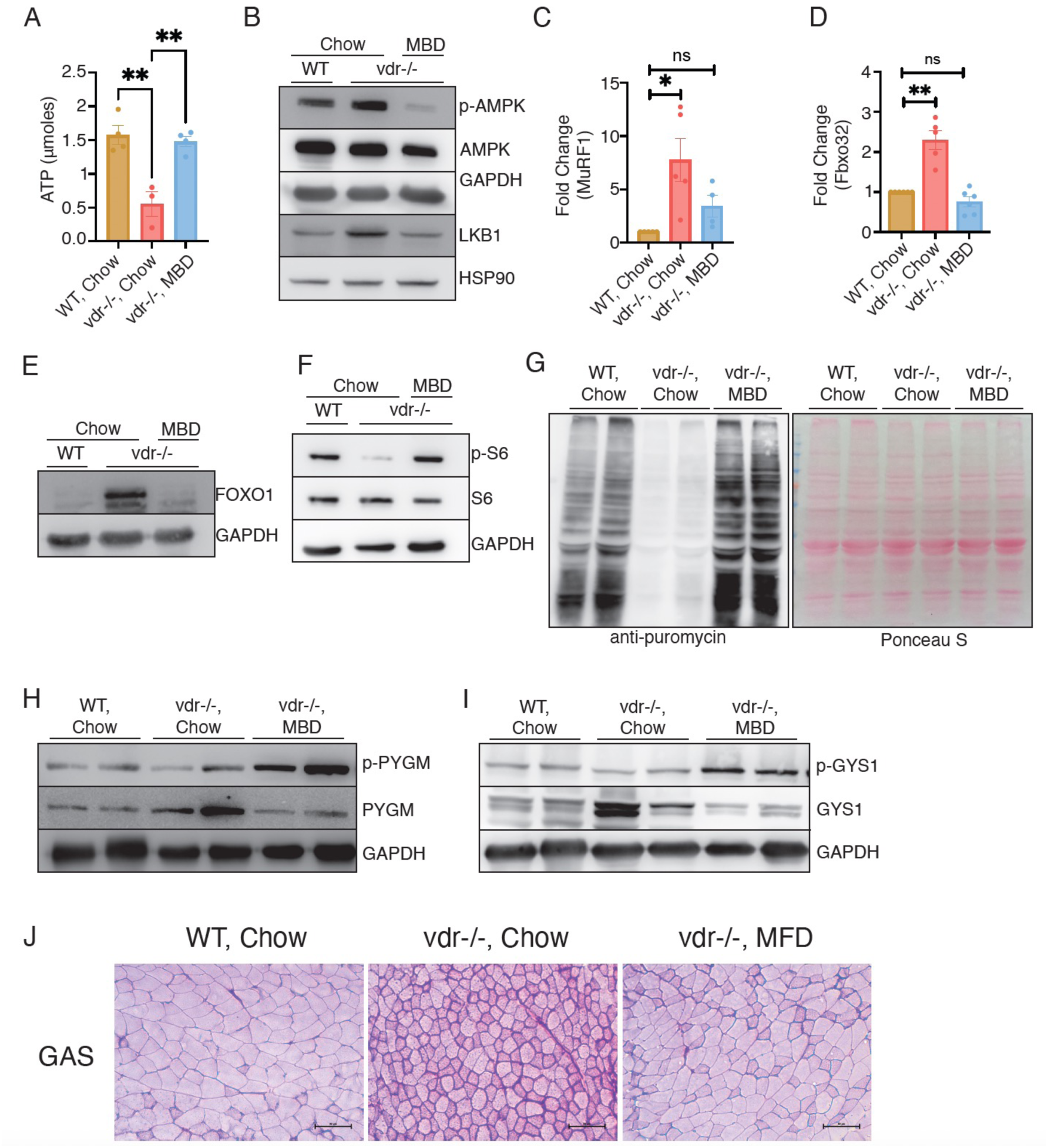
Milk-fat enriched diet restores the protein and energy homeostasis in skeletal muscles. **A.** ATP levels estimated in skeletal muscles of WT and vdr−/− mice in chow and MBD diets. **B.** Representative western blot for p-AMPK, AMPK and LKB-1 protein levels in hindlimb skeletal muscles of 7-week-old WT and vdr-/- mice on chow and MBD diets. **C-D.** Expression levels of Fbxo32 (C) and MuRF1 (D) mRNAs in QUAD muscles of 7-week- old WT and vdr−/− on chow and MBD diets quantified using qRT–PCR. **E.** Representative western blot for FoxO1 protein levels in hindlimb of skeletal muscles of 7- week-old WT and vdr-/- mice on chow and MBD diets (n=4). **F.** Representative western blot for p-S6 and S6 protein levels in hindlimb of skeletal muscles of 7-week-old WT and vdr-/- mice on chow and MBD diets (n=4). **G.** Western blot using anti-puromycin antibody for estimation of fasting protein synthesis in 7- week-old WT and vdr-/- mice on chow and MBD diets. Image of a Ponceau stained blot is used as loading control. Quantification of the blots are represented in supplementary figure 3C. **H-I.** Representative western blots showing protein levels of p-PYGM and PYGM (H) and p- GYS1 and GYS1 (I) in skeletal muscles of 7-week-old WT and vdr-/- mice on chow and MBD diets (n=4). **J.** Micrographs of PAS stained transverse sections of GAS muscles from WT and vdr−/− mice on chow and MFD diets. 10 μm cryosections of gastrocnemius muscles were used for staining. Magnification: 20X. Scale bars indicate 50μm. All graphs show mean ± SEM. *p < 0.05, **p < 0.01, ***p < 0.001, ****p < 0.0001 by one-way ANOVA (A, C, and D). Number of samples are denoted by the dots in the graphs.

Deregulation of protein homeostasis is associated with most occurrences of muscle atrophy. So, we further investigated the effect of milk-based diets on muscle proteostasis. MuRF1 and Fbxo32 are muscle-specific ubiquitin ligases upregulated during most of the known muscle wasting conditions, making them bona fide markers for muscle atrophy. We observed that vdr-/- on chow exhibited increased expression of these ubiquitin ligases. However, they were reduced to WT levels in the MBD-fed group in accordance with the alleviation of atrophy in the milk-like diets (Figures 3C & D). FOXO factors are stress-induced transcription factors well established to activate protein degradation pathways during atrophy. We have previously shown that Foxo1 is dramatically upregulated in the skeletal muscles of vdr-/- mice on chow (Das, Gopinath, and Arimbasseri 2022). Interestingly, milk-based diets downregulated Foxo1 protein expression in vdr-/- muscles (Figure 3E & Supplementary Figure 3B). These results indicate that milk-based diets alleviate both the energy stress and restores protein homeostasis of these muscles.

Next, we assessed if milk-based diets could alter protein synthesis in vdr-/- muscles. To address this, we first checked the activity of the mTORC1 pathway, an intracellular nutrient sensor essential for the upregulation of protein synthesis and muscle mass (Yoon 2017). As expected, vdr-/- mice on chow exhibited reduced phosphorylation of ribosomal protein S6, a downstream target of the mTORC1 pathway. The milk-based diet restored the phospho-S6 levels in these muscles, indicating the upregulation of the mTORC1 pathway (Figure 3F). Our previous work had shown that skeletal muscles of vdr-/- mice exhibit reduced fasting protein synthesis compared with WT mice (Das, Gopinath, and Arimbasseri 2022). We further analyzed if milk-based diets could restore the rate of fasting protein synthesis in vdr-/- muscles. We performed in vivo SunSET assay where puromycin incorporation in nascent proteins 30 minutes after intraperitoneal injection was assayed by western blot with an anti-puromycin antibody (Goodman and Hornberger 2013). As observed earlier, we found lower fasting protein synthesis in vdr-/- mice on chow. However, MBD-fed mice exhibited higher puromycin incorporation (Figure 3G), suggesting that the defect in protein synthesis of vdr-/- is corrected by MBD. Interestingly, the puromycin incorporation in vdr-/- on MBD was significantly higher than WT (Supplementary Figure 3C). These results imply that nutritional and metabolic defects in the vdr-/- skeletal muscles can be corrected by milk-based diets.

The skeletal muscles of vdr-/- mice exhibit a glycogen storage disorder characterized by increased glycogen synthase (GYS1) activity and reduced glycogen phosphorylase (PYGM) activity (Das, Gopinath, and Arimbasseri 2022). So, we asked if the milk-based diets could alleviate the glycogen storage disorder in vdr-/- mice. Western blot analyses show an increase in the activating phosphorylation of PYGM at ser15 and inhibitory phosphorylation of GYS1 at ser641 in vdr-/- mice on MBD (Figure 3H & I). Accordingly, PAS staining shows a reduction in glycogen accumulation in the GAS muscles of vdr-/- on MBD (Figure 3J). Taken together, these results indicate that milk based diets could restore muscle energy metabolism, muscle mass and growth of vdr-/- mice.

### The milk-based diet increases mitochondrial activity and lipid metabolism in the skeletal muscles of vdr-/- mice

We hypothesized that the milk-based diets alleviate muscle energy deprivation and atrophy by shifting muscles to fatty acid oxidation as the primary energy source. High fat-containing diets shift skeletal muscle metabolism towards fatty acid oxidation, sparing glucose as the primary energy source (P J Randle, Newsholme, and Garland 1964). Our RNAseq data also shows increased expression of genes involved in fatty acid oxidation (Figure 1). Free fatty acids are the preferred energy source for muscle when subjected to high-fat diets and under starvation conditions (Philip J. Randle 1998). Indeed, serum non-esterified fatty acid (NEFA) levels were higher in vdr-/- on MBD than in other groups (Figure 4A). In support of metabolic shift in vdr-/- muscles, we found that the expression of PDK4 protein, a kinase that inhibits PDH and thereby reorients the energy metabolism towards fatty acid oxidation (Sugden et al. 2000; Zhang et al. 2014), is upregulated in vdr-/- on both chow and MBD. Moreover, BODIPY-493/503 staining of the GAS muscle sections shows increased lipid droplets in vdr-/- on MBD (Figure 4C & D).

**Figure 4:**
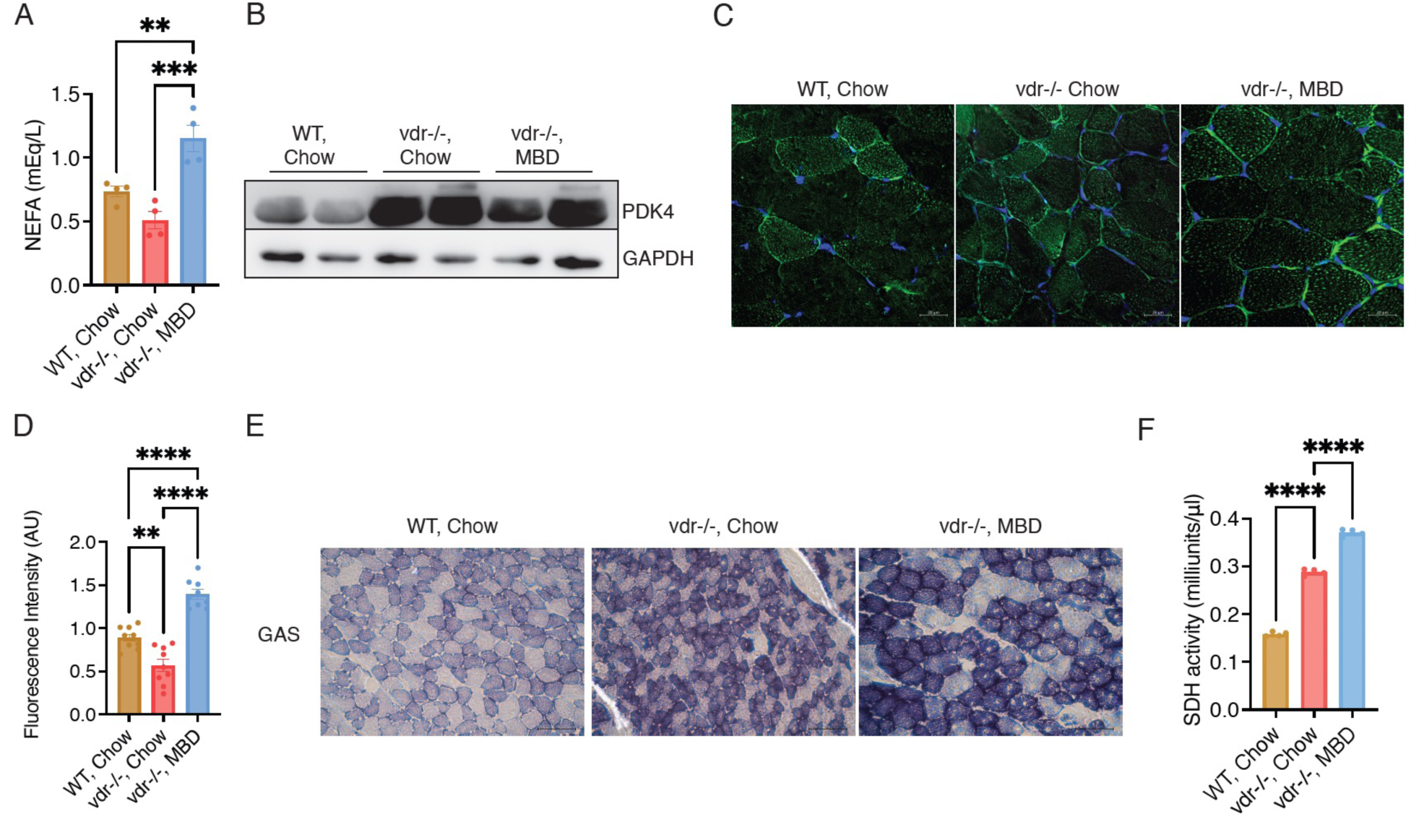
The milk-based diet increases mitochondrial activity and lipid metabolism in the skeletal muscles of vdr-/- mice. **A.** Serum non-esterified fatty acid (NEFA) levels in WT and vdr−/− mice on chow and MBD diets. **B.** Representative western blots showing protein levels of PDK4 in skeletal muscles of 7-week- old WT and vdr-/- mice on chow and MBD diets (n=3). **C-D.** Representative image of Bodipy staining of lipid droplets in the skeletal muscle (GAS) sections of WT and vdr-/- mice fed on chow and MBD diets (n=4). Quantifications of the same are represented in D. **E.** 10 μm cryo-sections of WT and vdr-/- skeletal muscle (GAS) sections of mice fed on chow and MBD diets stained with SDH buffer to estimate enzymatic activity in muscle sections. Magnification: 20X, scale: 10um. **F.** Succinate dehydrogenase complex (complex II) activity measured in gastrocnemius muscles of WT and vdr-/- mice fed on chow and MBD diets (n=4). All graphs show mean ± SEM. *p < 0.05, **p < 0.01, ***p < 0.001, ****p < 0.0001 by one-way ANOVA (A, D, and F). Number of samples are denoted by the dots in the graphs.

Since fatty acid catabolism requires mitochondrial activity, we checked if milk-based diets affect the mitochondrial abundance or activity in vdr-/- mice. Quantifying the mitochondrial DNA in the skeletal muscles of WT and vdr-/- mice showed no significant differences in chow or MBD (Supplementary Figure 4A). There was no statistically significant difference in the levels of various mitochondrial proteins involved in oxidative phosphorylation, such as SDHA (Complex II), UQCRC1 (complex III), and COX4 (complex IV) (Supplementary Figure 4B-D). To check if mitochondrial dynamics is altered in these mice, we investigated the expression levels of mitochondrial fusion proteins in the inner and outer mitochondrial membrane (OPA1 and MFN2). However, we did not observe any alteration in their expression (Supplementary Figure 4E & F).

To further confirm if vdr-/- mice on MBD exhibit altered mitochondrial activity, we assayed SDH (complex II) activity in cryosections of GAS muscles. As shown previously, SDH activity was higher in vdr-/- on chow than WT, and it increased further when vdr-/- mice were subjected to MBD (Figure 4E). SDH estimation in GAS muscle extracts confirmed the increased SDH activity observed in cryosections (Figure 4F), indicating increased mitochondrial activity in vdr-/- mice upon feeding MBD. These results reiterate that vdr-/- skeletal muscles do not exhibit defective mitochondrial biogenesis. Moreover, these strongly suggest that these muscles rely primarily on mitochondrial activity to utilize fat as the primary energy source. This supports the hypothesis that the inability of the vdr-/- muscles to efficiently utilize glucose primes them to shift to FAO and oxidative phosphorylation as the primary mechanism to generate energy and fat-rich diets indeed can restore muscle mass, energy, and growth.

### A lard-based, high-fat diet restores muscle energy levels but fails to fully restore muscle mass

Next, we asked if the ability to restore muscle mass and energy levels is specific to milk-based diets or if any fat source could restore the growth and muscle mass of the vdr-/- mice. To address this, we subjected these mice to a high-fat diet (HFD) with similar fat levels as MFD, while the source of fat was lard rather than butter (Supplementary Table 2). Both body size and weight increased when vdr-/- were subjected to HFD (Figure 5A). H&E-stained muscle sections show increased muscle fiber cross-section area in vdr-/- on HFD compared with chow (Figure 5B). However, the increase in cross-section area in HFD is intermediate compared with MBD (Supplementary Figure 5A). Unlike the milk-based diets, wet weights of TA, GAS, and QUAD muscles were also not restored to that of WT in these mice (Figures 5C & D and Supplementary Figure 5B). These results indicate that HFD was less effective in restoring muscle mass than milk-based diets.

**Figure 5:**
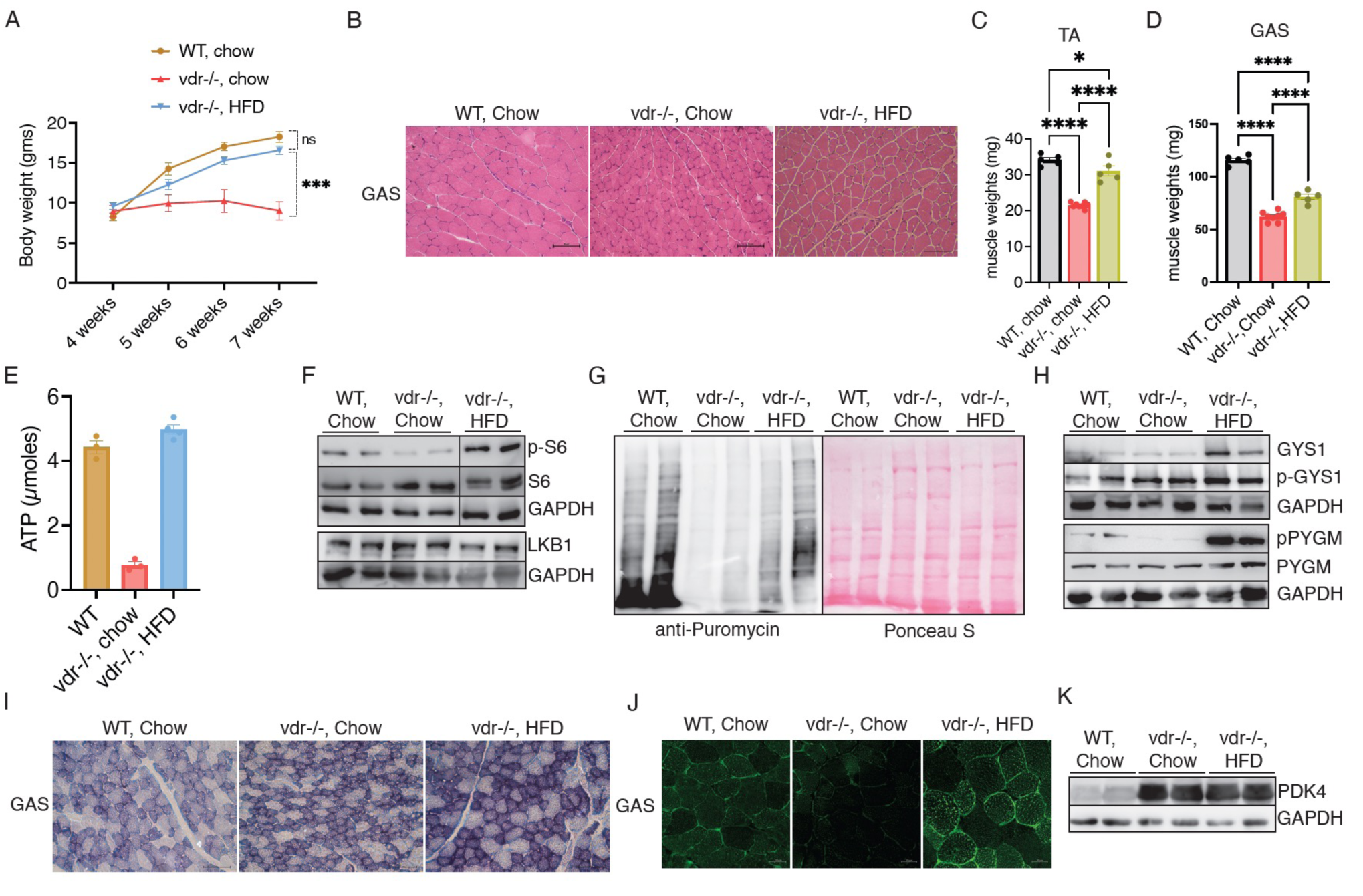
A lard-based, high-fat diet restores muscle energy levels but fails to fully restore muscle mass. **A.** Body weight (in grams) of WT and vdr−/− mice at 4 to 7 weeks of age on Chow and HFD diets (*n* > 6). **B.** H&E stained transverse sections of gastrocnemius (GAS) muscles of WT and vdr−/− mice at 7 weeks of age on Chow and HFD diets (n > 5). **C-D.** Tibialis anterior (TA) (C) and gastrocnemius (GAS) (D) muscle weight of 7-week-old WT and vdr−/− on Chow and HFD diets. **E.** ATP levels estimated in skeletal muscles of WT and vdr−/− mice in chow and HFD diets. **F.** Representative western blot for pS6, S6 and LKB1 protein levels in hindlimb of skeletal muscles of 7-week-old WT and vdr-/- mice on chow and HFD diets. **G.** Western blot using anti-puromycin antibody for estimation of fasting protein synthesis in 7- week-old WT and vdr-/- mice on chow and HFD diets. Image of a Ponceau stained blot is used as loading control. **H.** Representative western blots showing protein levels of p-GYS1, GYS1 and p-PYGM, PYGM in skeletal muscles of 7-week-old WT and vdr-/- mice on chow and HFD diets. **I.** 10 μm cryo-sections of WT and vdr-/- skeletal muscle (GAS) sections of mice fed on chow and HFD diets stained with SDH buffer to estimate enzymatic activity in muscle sections. Magnification: 20X, scale bar = 10um. **J.** Bodipy staining of lipid droplets in the skeletal muscle (GAS) sections of WT and vdr-/- mice fed on chow and HFD diets. **K.** Representative western blots showing protein levels of PDK4 in skeletal muscles of 7-week- old WT and vdr-/- mice on chow and HFD diets. All graphs show mean ± SEM. *p < 0.05, **p < 0.01, ***p < 0.001, ****p < 0.0001 by two-way ANOVA (A) or one-way-ANOVA (C, D, and E). Number of samples are denoted by the dots in the graphs.

Next, we asked if these muscles exhibit any defect in the muscle energy metabolism as in the case of vdr-/- on chow. Interestingly, muscle ATP levels were restored to WT levels in vdr-/- on HFD (Figure 5E). Accordingly, the energy-sensing AMPK pathway is downregulated in HFD fed group (Figure 5F). Moreover, the nutrient-sensing mTORC1 was also restored to WT levels in HFD-fed mice, indicating that HFD alleviated the energy deficiency in vdr-/- mice (Figure 5F). We further analyzed the rate of protein synthesis in these mice by SunSET assay. As predicted by the restoration of mTORC1 and energy levels in these mice, we found that the protein synthesis levels were upregulated in vdr-/- on HFD compared with chow (Figure 5G). However, the quantification of the signals indicate that the protein synthesis levels are not restored to that of WT level (Supplementary Figure 5C). By contrast, MBD exhibited significantly higher puromycin incorporation than even WT (Supplementary Figure 3C).

We further analyzed the glycogen metabolic pathways and storage in the skeletal muscles of WT and vdr-/- mice subjected to HFD. As in the case of MBD, we observed an increase in inhibitory phosphorylation at Ser641 of glycogen synthase (GYS1) and activating Ser15 phosphorylation of glycogen phosphorylase (PYGM), suggesting that the HFD could alleviate the defective glycogen storage in vdr-/- (Figure 5H). Accordingly, PAS staining of the GAS muscles shows reduced glycogen storage in vdr-/- on HFD (Supplementary Figure 5D). These results indicate that, even though HFD could not restore muscle mass to that of WT levels, it could circumvent the energy deficiency caused by glycogen storage disorder in vdr-/- mice.

We checked if the inability to restore muscle mass by HFD is due to functional defects in mitochondria. There was no significant change in mitochondrial DNA levels in the muscles of vdr-/- on HFD (Supplementary Figure 5E). Further, we confirmed that the SDH activity was high in HFD-fed mice by histological staining (Figure 5I) and SDH estimation in muscle lysates (Supplementary Figure 5F). These results indicate that HFD did not reduce mitochondrial activity. Moreover, HFD also increased lipid content in the skeletal muscles (Figure 5J) and increased PDK4 activity (Figure 5K), as observed in MBD, indicating that these muscles primarily use lipids as the energy source. These results suggest that high-fat-containing diets could ameliorate the energy deficiency observed in the vdr-/- mice, irrespective of the fat source. However, only milk-based diets could fully restore muscle weight.

### High-fat diet causes defective glucose clearance

vdr-/- mice exhibit defective insulin response and glucose homeostasis, processes that are known to impact muscle mass. On chow diet, they exhibit hypoglycemia despite having a muted insulin response (Das, Gopinath, and Arimbasseri 2022). Furthermore, high-fat-containing diets are known to negatively affect glucose homeostasis. Defective glucose homeostasis in conditions such as diabetes are known to negatively affect muscle mass. So, we asked if the inability of HFD to completely restore muscle mass reflects any systemic defects in glucose homeostasis.

We found that both milk-based diets restored the fasting glucose levels in the vdr-/- mice (Figure 6A). Decreased serum lactate levels in MFD also support the restoration of systemic energy homeostasis by milk-based diets (Figure 6B). Next, we performed a glucose tolerance test to estimate the kinetics of glucose clearance in these mice. As shown previously, vdr-/- mice exhibited faster glucose clearance, which is aided by the upregulation of Glut1 expression (Figure 6C). Interestingly, mice subjected to both the milk-based diets exhibited glucose clearance equivalent to that of WT mice on chow. This indicates that these diets normalized the glucose homeostasis in vdr-/- mice (Figure 6C). Like milk-based diets, HFD also restored the fasting blood sugar in vdr-/- mice (Figure 6D). However, HFD-fed mice exhibit a severe delay in glucose clearance, clearly showing defective glucose homeostasis (Figure 6E).

**Figure 6:**
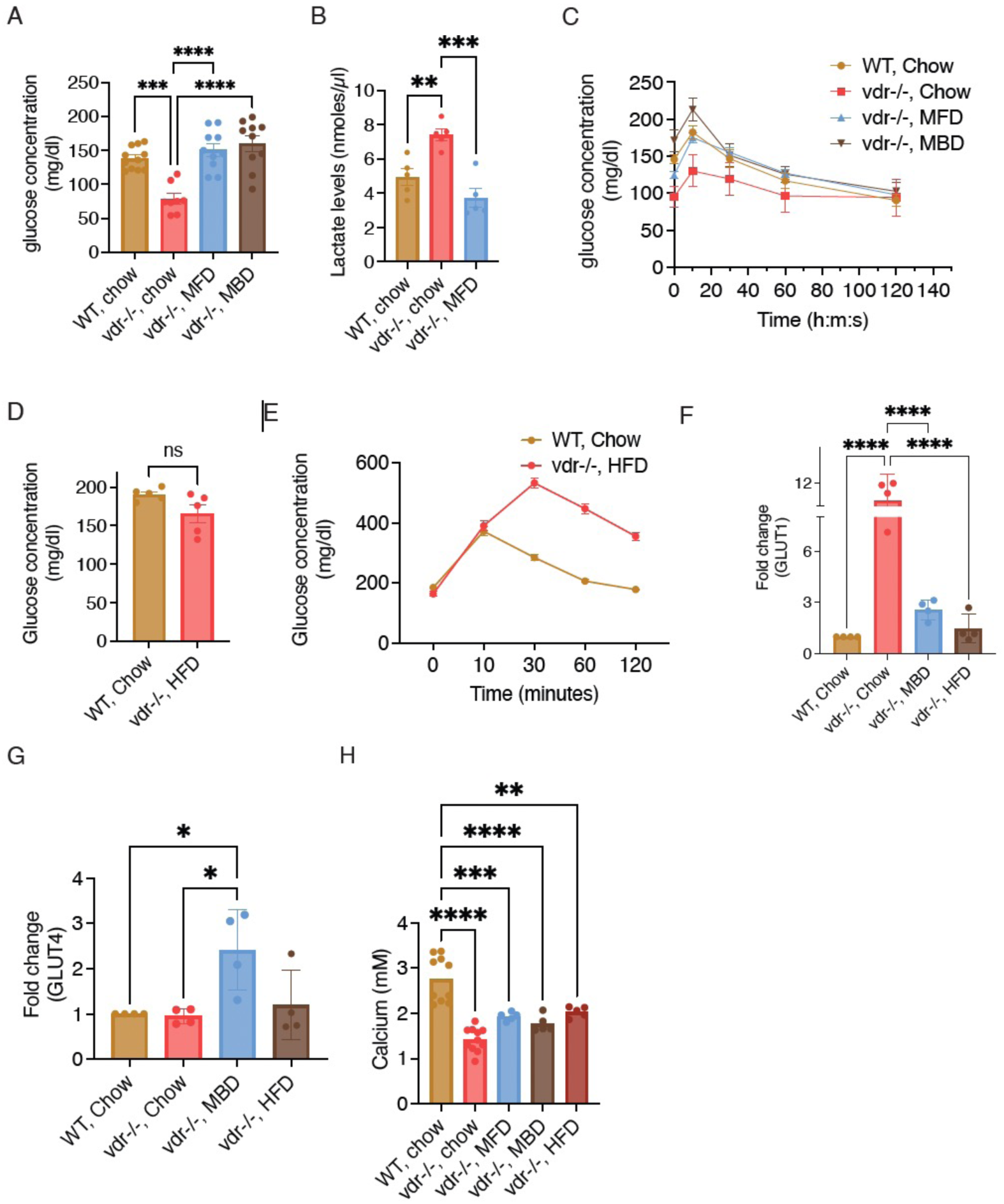
HFD causes defective glucose clearance. **A.** Fasting glucose levels of 7-week-old WT and vdr-/- mice fed on chow, MBD and MFD diets. **B.** Serum lactate levels in WT and vdr−/− fed on chow and MFD diets. **C.** Glucose tolerance test in 7-week-old WT and vdr-/- mice fed on chow, MBD and MFD diets (n ≥ 6). **D.** Fasting glucose levels of 7-week-old WT and vdr-/- mice fed on chow and HFD diets. **E.** Glucose tolerance test in 7-week-old WT and vdr-/- mice fed on chow, and HFD diets (n ≥ 6). **F-G.** qRT–PCR quantitation of GLUT1 (A) and GLUT4 (B) in WT and vdr−/− mice fed on chow, MBD and HFD diets. **H.** Serum calcium levels in WT and vdr−/− mice on chow, MBD, MFD and HFD diets. All graphs show mean ± SEM. *p < 0.05, **p < 0.01, ***p < 0.001, ****p < 0.0001 by one-way ANOVA (A, B, C, D, and H) or unpaired t-test (F). Number of samples are denoted by the dots in the graphs.

Skeletal muscles are the major sites of glucose deposition, which employs both insulin-dependent and insulin-independent transporters. AMPK pathway activation in vdr-/- mice on chow enables increased expression of GLUT1 and faster glucose uptake by muscles despite having no insulin response (Das, Gopinath, and Arimbasseri 2022). So, we checked if the differences in the glucose clearance observed between milk-based diets and HFD were associated with muscle glucose transporter expression. Both MBD and HFD lowered GLUT1 mRNA levels to WT levels consistent with the downregulation of AMPK pathway (Figure 6F). However, the transcript levels of GLUT4, the main insulin-responsive glucose transporter, increased in MBD treated mice but not in HFD fed ones (Figure 6G), indicating that the glucose clearance defect of HFD fed mice is due to reduction in expression levels of both Glut1 and Glut4, while MBD fed mice may be compensating for reduced Glut1 expression by upregulating Glut4 expression.

Taken together these results indicate that even though HFD could restore muscle energy levels by circumventing the defective carbohydrate utilization pathway, it could not restore muscle mass. Moreover, it led to further defects in systemic glucose homeostasis. On the other hand, milk-based diets could restore muscle energy metabolism, muscle mass, and systemic glucose homeostasis.

Low serum calcium levels are one of the major contributors to the drastic phenotype observed in vdr-/- mice (Endo et al. 2003), and milk is known to enhance calcium absorption independent of vitamin D <u>(Song, Kato, and Fleet 2003)</u>. So, we asked whether increased serum calcium levels bring about the differential effects of milk-based and lard-based high-fat diets. As expected, serum calcium levels were low in vdr-/- mice fed with chow. Interestingly, neither milk-based diets nor HFD could restore the serum calcium levels in vdr-/-, clearly suggesting that the positive effects of milk-based diets on muscle mass and metabolism are not due to increased serum calcium levels (Figure 6H).

### Milk-based diet could restore insulin response in vdr-/- but high fat diet led to inhibition of insulin synthesis

Insulin is the essential hormone for maintaining glucose homeostasis and muscle mass. Insulin-stimulated glucose uptake and upregulation of anabolic pathways are essential for maintaining skeletal muscle mass. Insulin secretion is defective in mice lacking functional VDR (Zeitz et al. 2003). Since we observed a dichotomy between milk-based diets and HFD regarding glucose clearance and expression levels of insulin sensitive glucose transporter Glut4 expression, we asked if there is any difference in insulin response between these two groups. Indeed, vdr-/- on chow exhibited deficient insulin levels in both fasted and fed states (Figure 7A). On the other hand, MBD-fed mice show fasting and fed state insulin levels similar to WT mice indicating that MBD can restore pancreatic insulin production/secretion. In contrast, HFD fed vdr-/- mice exhibit very low insulin levels in the fasting state. Though insulin levels increased upon feeding, indicating a response to glucose, its levels were still very low, comparable with fasted WT mice (Figure 7A). These results suggest that increased insulin levels together with higher Glut4 levels restore glucose clearance in vdr-/- mice subjected to MBD, while the low insulin levels in HFD could be responsible for delayed glucose clearance and lower muscle mass in that group.

**Figure 7:**
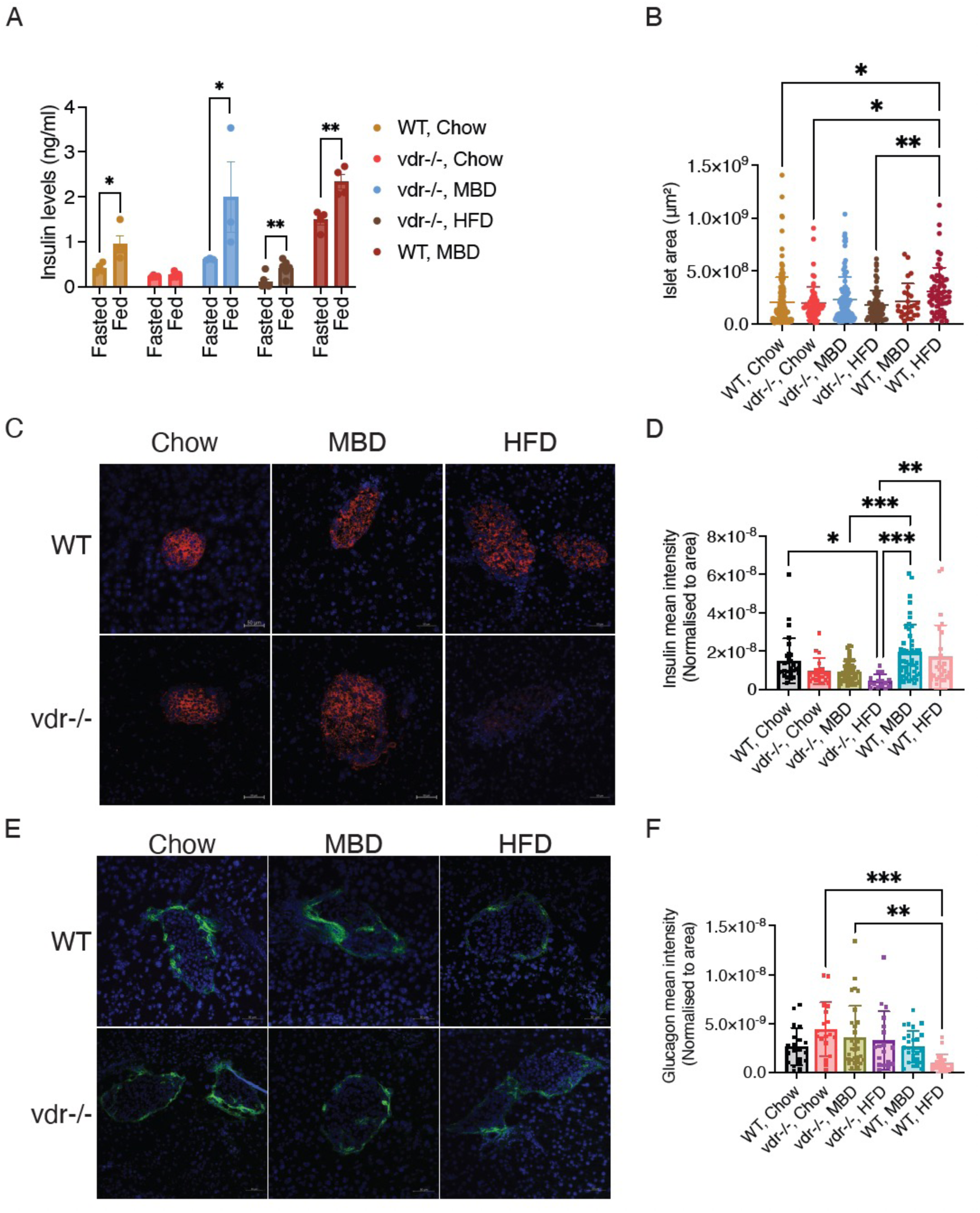
Milk-based diet could restore insulin response in vdr-/- but HFD led to inhibition of insulin synthesis. **A.** Serum insulin levels measured by ELISA at fasting state and 10 min after intraperitoneal injection of glucose in WT and vdr−/− mice fed on chow, MBD and HFD diets. **B.** Islet area in 7-week-old WT and vdr-/- mice fed on chow, MBD, and HFD diets. **C,E.** Confocal microscopy analysis of insulin and glucagon staining performed on islets from 7 weeks old WT and vdr-/- mice fed on chow, MBD, and HFD diets. Scale bar = 50 um. **D,F:** Mean fluorescence intensity measurements of insulin (C) and glucagon (E) of islets from 7 weeks old WT and vdr-/- mice fed on chow, MBD, and HFD diets. All graphs show mean ± SEM. *p < 0.05, **p < 0.01, ***p < 0.001, ****p < 0.0001 by paired t- test (A) or one-way ANOVA (B, D, and F). Number of samples are denoted by the dots in the graphs.

Further analysis of pancreas shows that vdr-/- on chow exhibits reduced pancreas weight, but both the milk-based diet and HFD restored this to WT levels (Supplementary Figures 6A & B). However, normalization of the pancreas weight with body weight shows lower weight in HFD fed mice. Histopathology analysis of paraffin-embedded pancreas sections did not show any difference in islet area for vdr-/- on HFD (Figure 7B). However, intriguingly, WT on HFD shows an increase in islet area compared with vdr-/- on chow and HFD. Islet morphology was also normal in vdr-/- on HFD group (Supplementary Figure 6C). To test whether these diets affect insulin content in the pancreas, we performed immunofluorescence analysis of pancreatic islets using an anti-insulin antibody. Though vdr-/- on chow had lower serum insulin, we found no statistically significant difference in the fluorescence intensities (normalized for the area) between WT mice on chow and vdr-/- on chow (Figure 7C & D), consistent with previous report that their defect in insulin response was due to reduced insulin secretion (Zeitz et al. 2003). MBD fed vdr-/- also exhibited signal intensity similar to WT on chow. However, vdr-/- on HFD shows a drastic decrease in the fluorescence intensity, indicating lower insulin content. These results suggest that HFD inhibited insulin synthesis in vdr-/- mice. On the other hand, WT mice on HFD did not show any decrease in insulin signals, clearly indicating that vitamin D signaling is essential for insulin synthesis by pancreatic beta cells when mice are subjected to a lard-based high-fat diet. To test whether this effect is limited to insulin or other hormones secreted by pancreatic islets are also affected, we analyzed glucagon levels in these mice. No change in glucagon signal intensity was observed in any groups except for WT on HFD (Figure 7E & F). Taken together, these results show that milk-based diets restore skeletal muscle energy metabolism and pancreatic insulin response in vdr-/- mice. On the other hand, lard based high-fat diet could not restore systemic energy metabolism, most likely due to defective insulin synthesis in pancreas.

## DISCUSSION

Metabolic flexibility of an organism depends on its ability to modulate biochemical reactions to match different energy sources (Smith et al. 2018). These are specifically important when major shifts in the feeding habits of the organism occurs. Weaning is a process in mammals when the organism makes such a shift from a fat-enriched milk diet to carbohydrate based diet, which most of the mammals follow throughout their life (Ferré et al. 1986). Previous studies have shown that vdr-/- mice delays the onset of growth defects and atrophy could be achieved by deferral of weaning (Endo et al. 2003). Since a VDR independent pathway maintains the serum calcium levels during the suckling stage, the delayed disease onset was attributed to the increased calcium levels (Song, Kato, and Fleet 2003). The results presented here shows that continuing these mice on a milk-based high fat diet alleviates many defects exhibited by vdr-/-. However, this reversal is independent of restoration of serum calcium levels and therefore represent an alternative metabolic state with limited dietary flexibility.

The role of vitamin D in regulation of systemic metabolism is increasingly appreciated. A comprehensive model that integrates the role of vitamin D in systemic metabolism is essential to assess the damages caused by vitamin D deficiency on metabolic flexibility. An increased metabolic rate and lean phenotype was observed in mice vdr-/- and Cyp27b1-/- mice (Su et al. 2021; Narvaez et al. 2009). Moreover, in male mice, vitamin D deficiency attenuates hyperinsulinemia and lipid accumulation in the liver (Liu et al. 2015). But the molecular mechanisms that lead to the hyper-metabolic rate in vdr-/- are not known. Cyp27b1 null mice exhibit upregulation of the renin angiotensin system and associated increase in corticotropin releasing hormone (CRH), which could be responsible for increased metabolic rate and lean phenotype (Su et al. 2021). On the other hand, in vdr-/- mice, upregulation of the uncoupling protein UCP1 in the adipose tissue correlated with their resistance to body weight gain even under high-fat diet (Narvaez et al. 2009). The evidence clearly indicates that vitamin D signaling deficiency indeed leads to a general energy depletion and this process involves multiple organs.

Our previous study had shown that skeletal muscles of vdr-/- mice exhibit increased storage and decreased utilization of glycogen (Das, Gopinath, and Arimbasseri 2022). In vdr-/- mice, the depletion of glucose as energy source due to its accumulation in muscles as glycogen, when combined with adipose tissue atrophy, could be leading to a systemic energy depletion (Zinser and Welsh 2004; Guzey et al. 2004; Narvaez et al. 2009). Feeding energy rich diets with higher fat content could indeed restore the skeletal muscle energy levels, supporting the hypothesis that muscle energy deficiency could be attenuated by changing the primary energy source. The restoration of muscle energy and muscle metabolism by MBD (38% of energy from fat) as well as MFD (60% energy from fat) reveals that even moderate increase in the fat levels is sufficient to make these changes. The upregulation of fatty acid oxidation genes and the mitochondrial activity of skeletal muscles of chow-fed vdr-/- mice suggested that these muscles have adapted to alternate energy sources as their carbohydrate utilization in impaired and provision of fat enriched diets could restore muscle metabolism and thereby systemic metabolism. Though we have not addressed it in this study, it appears that the high-fat diets could also restore adipose tissue depots in vdr-/- (data not shown).

However, the results obtained with the lard-based high-fat diet (60% energy from fat) contradicts this oversimplified model. This diet also led to an increase in skeletal muscle energy levels and signaling pathways such as the mTORC1 and the AMPK pathway but failed to restore the muscle weight to that of WT levels. This is consistent with their inability to upregulate fasting protein synthesis to the WT levels. The restoration of insulin response by the pancreas specifically by the milk-based diets could be an additional requirement to restore muscle mass and glucose clearance. Indeed the downregulation of Glut1 observed when the mice were fed on MBD and HFD could be due to downregulation of AMPK (Jing, Cheruvu, and Ismail-Beigi 2008). However, MBD could increase expression of Glut4, whose expression and localization are dependent on insulin signaling (Lu et al. 2015). But HFD did not show any difference in Glut4 levels. This distinction, together with the inability of the pancreas to generate insulin in the absence of vitamin D when subjected to HFD may have led to the lower muscle mass and defective glucose clearance in these mice. This model is summarized in Figure 8. Moreover, the decrease in skeletal muscle glycogen storage in vdr-/- mice fed with milk-based diets and HFD is consistent with the inhibition of glycogen synthesis by free fatty acids (G Boden and Chen 1995; Jaafar et al. 2019).

**Figure 8:**
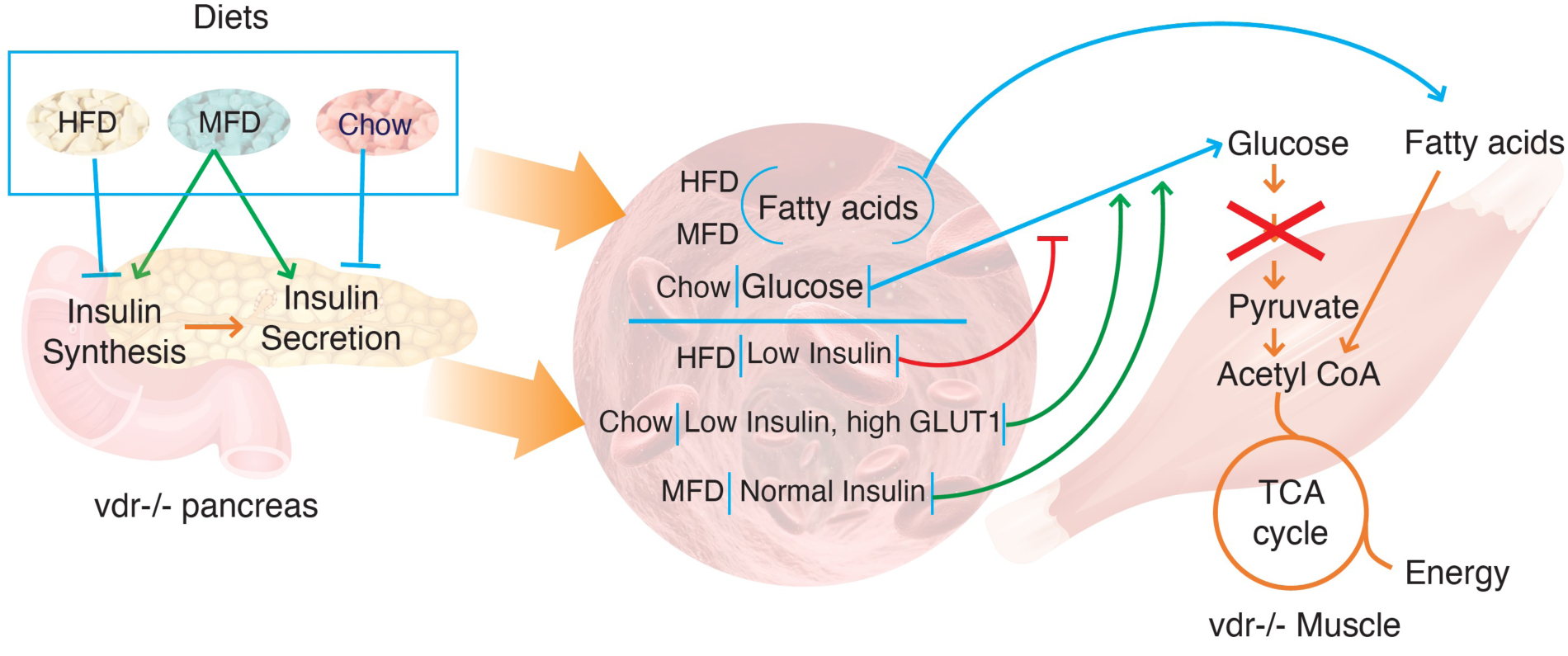
A model showing the interaction between pancreas and skeletal muscles under different dietary regimens. Pancreas: In the absence of VDR, chow diet leads to reduced insulin secretion and HFD leads to defective insulin synthesis. On the other hand, MBD normalizes the insulin synthesis and secretion. As a result, insulin levels are decreased in the serum of both chow-fed and HFD-fed mice. Muscle: In chow diet, the muscles takes up glucose independent of insulin, aided by the upregulation of Glut1. On the other hand, in HFD fed mice, muscle energy is restored by fatty acid oxidation, suppressing AMPK pathway and Glut1 expression. Since there is no insulin response in these mice, glucose uptake is diminished leading to delayed glucose clearance. In case of MBD mice, fatty acids restore muscle energy levels leading to reduced Glut1 expression. However, their insulin response is restored, leading to insulin dependent glucose uptake and normal glucose clearance.

An interesting observation made in this study is the normalization of insulin response in vdr-/- mice when subjected to milk-based diets, which was reflected in the glucose tolerance tests as well. Vitamin D is known to have negative effects on both insulin secretion and insulin sensitivity (Alvarez and Ashraf 2010; Ayesha et al. 2001; Norman et al. 1980). Mice with VDR DNA binding domain deletions show drastic decrease in the serum insulin levels due to defective insulin secretion (Zeitz et al. 2003). In accordance with these reports, we also have observed reduced serum insulin levels during fasting and in response to glucose stimulation. However, the insulin levels in islets did not vary in vdr-/- mice fed on chow diet suggesting that they show a defect in insulin secretion rather than synthesis. The insulin secretion was normalized when these mice were fed on milk-based diet. Since the milk-based diets could not restore the serum calcium levels, we argue that the increase in insulin secretion when these mice were subjected to MBD was independent of serum calcium levels. Increased serum free fatty acids levels, which was a feature of vdr-/- mice on milk based diets have been shown to stimulate insulin secretion (Guenther Boden 2005; Cen, Sargsyan, and Bergsten 2016; Nolan et al. 2006; Poitout 2018). A previous study had shown that weaning mice on to a milk-fat diet maintained high mTOCR1 activity, which in turn maintained higher basal insulin secretion (Jaafar et al. 2019). Accordingly, when the WT mice were fed with MBD, we also observed an increase in the basal insulin levels, which further increased upon stimulation with glucose. Further studies are required to understand the molecules and/or pathways that are activated by MBD to increase insulin synthesis and secretion.

By contrast to the restoration of insulin secretion by vdr-/- on milk-based diets, the reduced insulin levels in the islets of vdr-/- mice on HFD was surprising. The insulin levels were similar even in the islets of WT mice fed with HFD. So, a drastic decrease in the insulin levels of HFD fed vdr-/- mice indicates that under these specific conditions, insulin synthesis depends on vitamin D signaling (Figure 8). Identifying the determinants of the VDR dependence of insulin production under different dietary conditions will be of very interest and importance.

In conclusion, the metabolic inflexibility in the mice lacking VDR stems from the predisposition of skeletal muscle to primarily depend on fatty acids for energy metabolism. However, it appears that this is only one of the several layers of the effects vitamin D has on systemic metabolism. For example, though the muscle energy levels are restored in vdr-/- on HFD, they have systemic glucose imbalance due to defective insulin synthesis. Thus, vitamin D is also required when the mice are fed on a lard-based high-fat diet. Furthermore, the effect of these dietary conditions on other vitamin D sensitive metabolic tissues such as adipose and liver and other endocrine systems will be essential to reach a holistic model of the effect of vitamin D on metabolic homeostasis. Such a model is required to address the disorders associated with metabolic inflexibility, which arise due to several reasons including lifestyle, childhood or adult malnutrition and ageing.

## Materials and Methods

### Animals

Vitamin D receptor null mutant mice were purchased from Jackson Laboratories (Stock No. 006133, B6.129S4-Vdrtm1Mbd lJ; Bar Harbor, ME, USA) and maintained at the Small Animal Facility at the National Institute of Immunology, New Delhi, India. Genotyping was performed on each litter once heterozygous mating pairings were established. From the same litter, vdr null and WT were chosen as the experimental and control groups. Ad libitum feedings of a commercial rodent chow, milk-based diet (Lactogen 2, Nestle), milk fat diet (Research Diets), and high fat diet (Research Diets) were administered to all mice. Since the parameters evaluated did not differ between male and female mice, both were used for the study. Experiments were performed on mice at 3, 5, or 7 weeks of age as indicated in the text. Mice were housed in a sterile facility in individually ventilated cages (IVC). Maintenance of animal strain, tissue collection post sacrifice and drug administration were performed in accordance with the guidelines of the Animal Ethics Committee at the National Institute of Immunology.

### RNA extraction

Quadriceps muscles were dissected and snap-frozen in liquid nitrogen and homogenized in RNAiso plus Reagent (TAKARA, 1 mL/100 mg tissue) to isolate total muscle RNA as per the manufacturer’s recommendations. 400 µl chloroform was added and centrifuged at 10,000 rpm for 10 min at room temperature to separate the three phases. The supernatant containing RNA was carefully transferred to a fresh RNase free microfuge tube and re-extracted by adding equal volume of chloroform to it. RNA was precipitated by the addition of 0.5 volumes of isopropanol followed by incubation at RT for 10 minutes. The sample was centrifuged at 13,000 rpm for 30 min at 4°C, washed with 70% ethanol and air dried. The RNA pellet was dissolved in 50µl RNase free DEPC water. Total RNA was quantified with a Multiskan SkyHigh microplate spectrophotometer (Thermo Scientific, Wilmington), and a ratio of 260 to 280 nm was used to determine the RNA quality. 500 ng of sample was run on a 1.5% formaldehyde-agarose gel to check integrity.

### Quantitative RT–PCR

First-strand cDNA was synthesized from total RNA using the PrimeScript First-Strand cDNA Synthesis Kit with PrimeScript Reverse Transcriptase according to the manufacturer’s protocols (TAKARA). To 1 ug of RNA, 2 µl of reaction mix containing 50 pmoles of oligodT and 10 µmoles of dNTPs was added. Total volume was made up to 10 µl with DEPC treated water. The mix was heated at 65 degrees for 5 minutes and immediately cooled on ice. To this mixture, 20units of RNase inhibitor, 200 units of PromeScript reverse transcriptase, and 4µl of 5X buffer was added and kept at 42 degrees for 45 minutes followed by 70 degrees for 15 minutes. Quantitative RT– PCR was performed using the QuantStudio-6 and -7 Flex Real-Time PCR Systems (Thermo Fisher Scientific) with SYBR® Premix Ex Taq (Tli RNase H Plus) by TAKARA. Each sample was amplified in triplicates using primers specific to genes of interest. The expression levels of each transcript were normalized to the housekeeping gene GAPDH. The primer sequences used are mentioned in Supplementary Table 3.

### RNA sequencing and data analysis

Isolated RNA was used for library preparation using Truseq stranded RNAseq library preparation kit (Illumina) and subjected to sequencing on the Illumina platform (Macrogen, Seoul, S. Korea). FastQC package (http://www.bioinformatics.babraham.ac.uk/projects/fastqc) was used for quality control of the reads. Adapter trimmed reads were mapped using HISAT2 (Kim, Langmead and Salzberg, 2015) to mouse genome from Ensemble. Htseq-count (Kim, Langmead and Salzberg, 2015)(Anders, Pyl and Huber, 2015) was used for calculating the number of reads mapping to each gene. Differential expression analysis was performed using DESeq2 (Mainz Mainz Press, 2016). For enrichment analysis, GSEA (Subramanian et al., 2005) and g:profiler (Reimand et al., 2007) tools were used.

### Preparation of whole cell lysates and immunoblotting

Protein extracts from the hindlimb muscles of mice were obtained by homogenizing muscles in lysis buffer (50 mM Tris–HCl pH 7.5, 150 mM NaCl, 5 mM EDTA, 1% NP-40, 0.5% sodium deoxycholate, 0.1% SDS) containing protease/phosphatase inhibitor cocktail (Cell Signaling Technology). Protein estimation was done by the BCA method. Equal amount of protein for each sample was resolved by SDS-PAGE (8%, 10%, 12%, and 15%) at a constant voltage of 120V using the Biorad Mini protean tetra system. The proteins were transferred onto the nitrocellulose membrane (BioRad and Himedia) at a constant voltage of 120V for 2 hours at 4°C. Post transfer, the membranes were blocked in 2.5% BSA (Himedia, India) dissolved in TBST (0.1% Tween 20 in Tris Buffered saline) for 60 minutes. Following this, the membranes were incubated with primary antibodies prepared in 2.5% BSA in 1X TBST solution, overnight at 4°C in shaking condition. Then, the membranes were washed using TBST, thrice for 5 minutes each and incubated with HRP conjugated secondary antibodies (1:10000) prepared in 2.5% BSA in 1X TBST solution for 60 minutes at room temperature. The membranes were washed with TBST (thrice for 5 minutes each) and developed using Supersignal West Pico Chemiluminescent Substrate (Thermo Scientific, Rockford, IL, USA). The bands were visualized through ImageQuant LAS500 series (GE Healthcare). List of antibodies used are given in Supplementary Table 4.

### In vivo estimation of muscle protein synthesis (SUnSET assay)

To perform SunSET assay, mice were starved for 7 hours followed by an intraperitoneal injection of 0.040 μmol/g puromycin dissolved in 100 μL of PBS. Tibialis anterior (TA) muscle was removed at precisely 30 minutes after injection and immediately frozen in liquid nitrogen for western blot analysis. To detect puromycin incorporation, a mouse IgG2a monoclonal anti-puromycin antibody (clone 12D10, 1:5000) was used.

### Glucose tolerance assay

To perform Glucose tolerance assay, WT and vdr -/- mice were starved for seven hours in their regular cage environment to investigate changes in glucose tolerance. The mice had unrestricted access to water during that period. At time 0, an intravenous injection of glucose (2 mg/kg body weight) was administered in saline. A small drop of blood from the tip of the tail was placed on to a test strip, and the Accu-Chek Active Glucometer was used to record the readings in order to determine the blood glucose levels in whole blood at 0, 10, 30, 60 and 120 min.

### Serum insulin and lactate estimation

From the same mice, serum samples were taken in order to estimate the levels of lactate and insulin. Serum was isolated from the retro orbital venous plexus of mice that had been fasted for seven hours at baseline and 30 minutes after an intraperitoneal glucose challenge. According to the manufacturer’s protocol, serum lactate was calculated using the lactate estimation kit (Abcam #ab65331). Serum insulin levels were determined by ELISA using mouse insulin ELISA kit from Sigma according to the manufacturer’s instructions.

### ATP estimation in muscles

ATP estimation was performed using the kit from Thermo Scientific according to manufacturer’s protocol. A modified RIPA buffer with EDTA (150 mM NaCl, 1% NP-40, 0.5% Na-deoxycholate, 0.1% SDS, 25 mM Tris, pH 7.4) was used to sonicate a total of 100 mg of tissue and the lysate was centrifuged at 13000 RPM for 15 min at 4°C. Post supernatant collection, the assay was carried out in accordance with the manufacturer’s instructions.

### Cryo-sectioning and histological analyses

Skeletal muscle (tibialis anterior and gastrocnemius) for histology were collected from 7-weekold mice and frozen in isopentane chilled in liquid nitrogen. 10µm sections were cut using cryotome (Thermo Scientific HM525 NX). These sections were stained with Hematoxylin & Eosin (H&E) and periodic acid-Schiff (PAS) according to the manufacturer’s protocol (Sigma). The muscle sections were analyzed blindly by a pathologist, and the images were obtained at ×400 magnification (scale bar = 20 μm). For succinate dehydrogenase (SDH) staining of muscle sections, frozen sections of skeletal muscles were incubated for 50 min in a humidified chamber at 37°C with incubation media [100 mM phosphate buffer, pH 7.6, 1.2 mM nitroblue tetrazolium (NBT), and 100 mM sodium succinate] followed by imaging. For BODIPY staining, muscle sections were incubated for 30 minutes at room temperature with BODIPY (Invitrogen), washed thrice with 1X PBS and mounted with DAPI containing mounting media (UltraCruz). The sections were imaged by confocal microscope (Zeiss LSM 980). For pancreatic histology, the pancreata were embedded in OCT compound (Thermo Scientific) and flash-frozen in isopentane chilled on liquid nitrogen. 7-μm thickness of the frozen tissues were sectioned, mounted on slides, and stored at −80 °C. For insulin and glucagon staining, the tissue sections were fixed in pre-cooled acetone for 10 minutes. The slides were rinsed twice with 1X PBS at a neutral pH for 5 minutes followed by blocking (10% fetal bovine serum in 1X PBS) at RT for an hour. For insulin immunohistochemistry, slides were incubated with rabbit insulin antibody (Insulin Antibody #4590, Cell signaling Technology) overnight at 4C, followed by incubation with Anti rabbit Alexa Fluor™ 568 (1/500) for 1 hour at RT. For glucagon staining, the same procedure was repeated with primary rabbit glucagon antibody (Glucagon Antibody #2760, Cell signaling Technology) and Goat anti-Rabbit IgG (H+L) Alexa Fluor™ Plus 488 as secondary antibody. Sections were imaged by confocal microscope (Zeiss LSM 980). The average pixel intensity of insulin and glucagon positive staining per islet was measured, following background subtraction, to determine the insulin and glucagon content in pancreatic islets. Using Image J software, all images were taken from randomly chosen pancreatic regions using the same parameters. Hematoxylin & Eosin (H&E) staining of pancreatic sections were performed according to the manufacturer’s protocol (Sigma). The islet area was quantified using the same using ImageJ software.

### Estimation of serum calcium, muscle succinate dehydrogenase and pyruvate dehydrogenase

Enzymatic Assays from skeletal muscle tissues were done from muscle lysates prepared in the assay buffer as per the manufacturer’s protocol (Sigma-Aldrich). Serum calcium levels were measured using calcium estimation kit (Abcam #ab102505) according to the manufacturer’s instructions.

### Serum lipid profiling

Sera samples were collected by retro-orbital bleeding in mice and used for this assay. Levels of cholesterol, triglyceride, High density lipoprotein (HDL), Low density lipoprotein (LDL), Urea, Alanine transaminase (ALT/SGPT) and Aspartate transaminase (AST/ SGOT) were measured using Coralyser Smart Coral Clinical Systems (Tulip Diagnostics Pvt Ltd).

### Statistical analysis

Western blots were densitometrically quantified using ImageJ software. All experiments were performed at least three times. Data points in the graphs represent the number of samples for each figure. Statistical tests performed for each assay varied based on the nature of comparison, and data distribution. The details of tests used in each figure is described in the figure legends. In general, if the comparison was between two groups, paired or unpaired t-tests were used. When there were more than two groups were compared, one-way or two-way ANOVA was used. To ascertain whether samples had equal variance or unequal variance, F test was used. Error bars denote the standard error of the mean (SEM), while graphs display the mean of the specified number of replicates. The GraphPad Prism software, version 9, was used to create each graph.

## Supporting information

Supplemental Data Set 1

---

**Supplementary figure 1:**
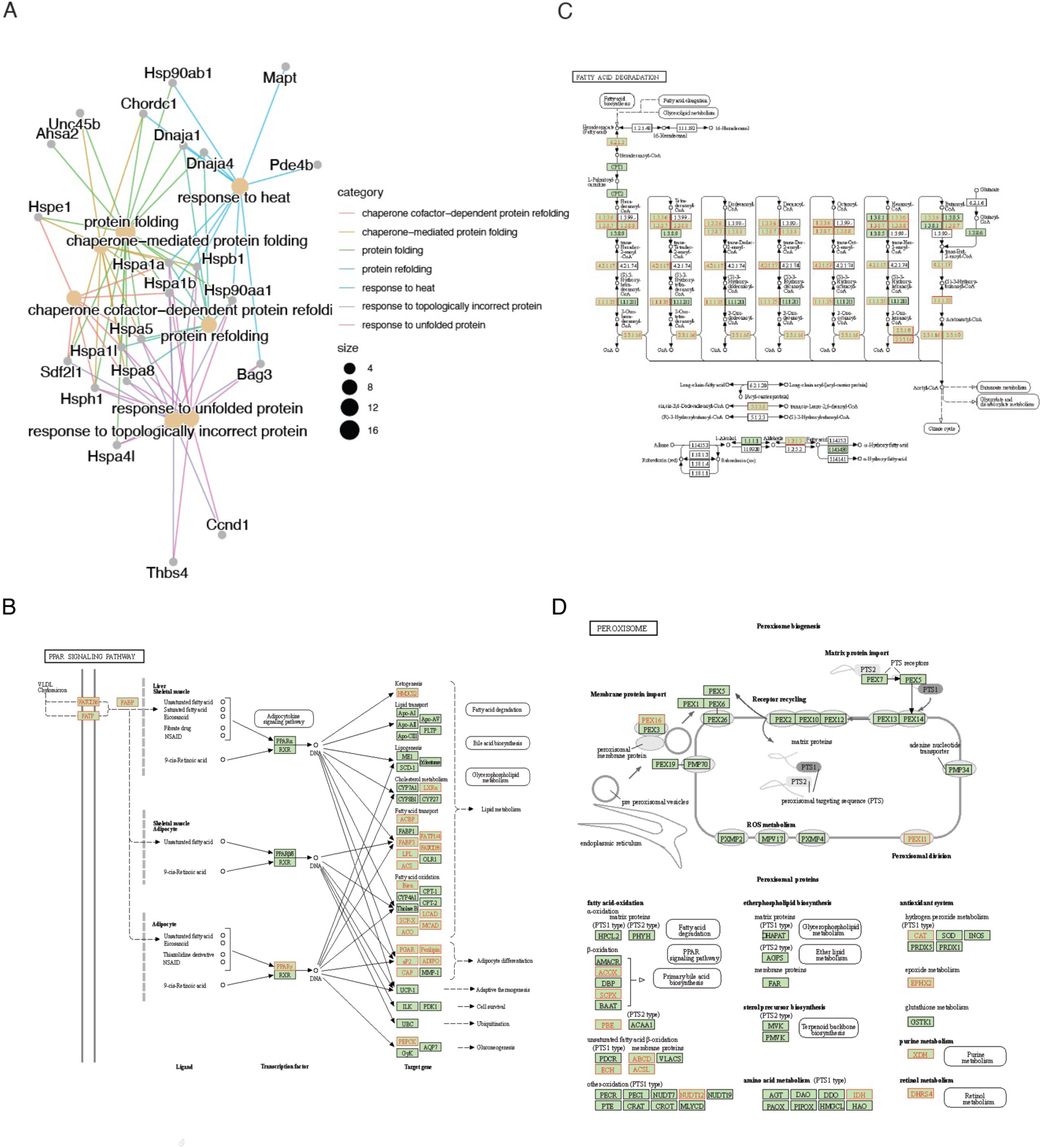
vdr-/- skeletal muscles are predisposed for fatty acid catabolism. **A.** A network plot showing downregulated genes in skeletal muscles of 7 weeks-old vdr-/- mice enriched in functions associated to protein folding. **B-D.** KEGG pathways PPAR signalling (B), fatty acid degradation (C) and peroxisome (D) shown with genes upregulated in vdr-/- highlighted in red.

**Supplementary figure 2:**
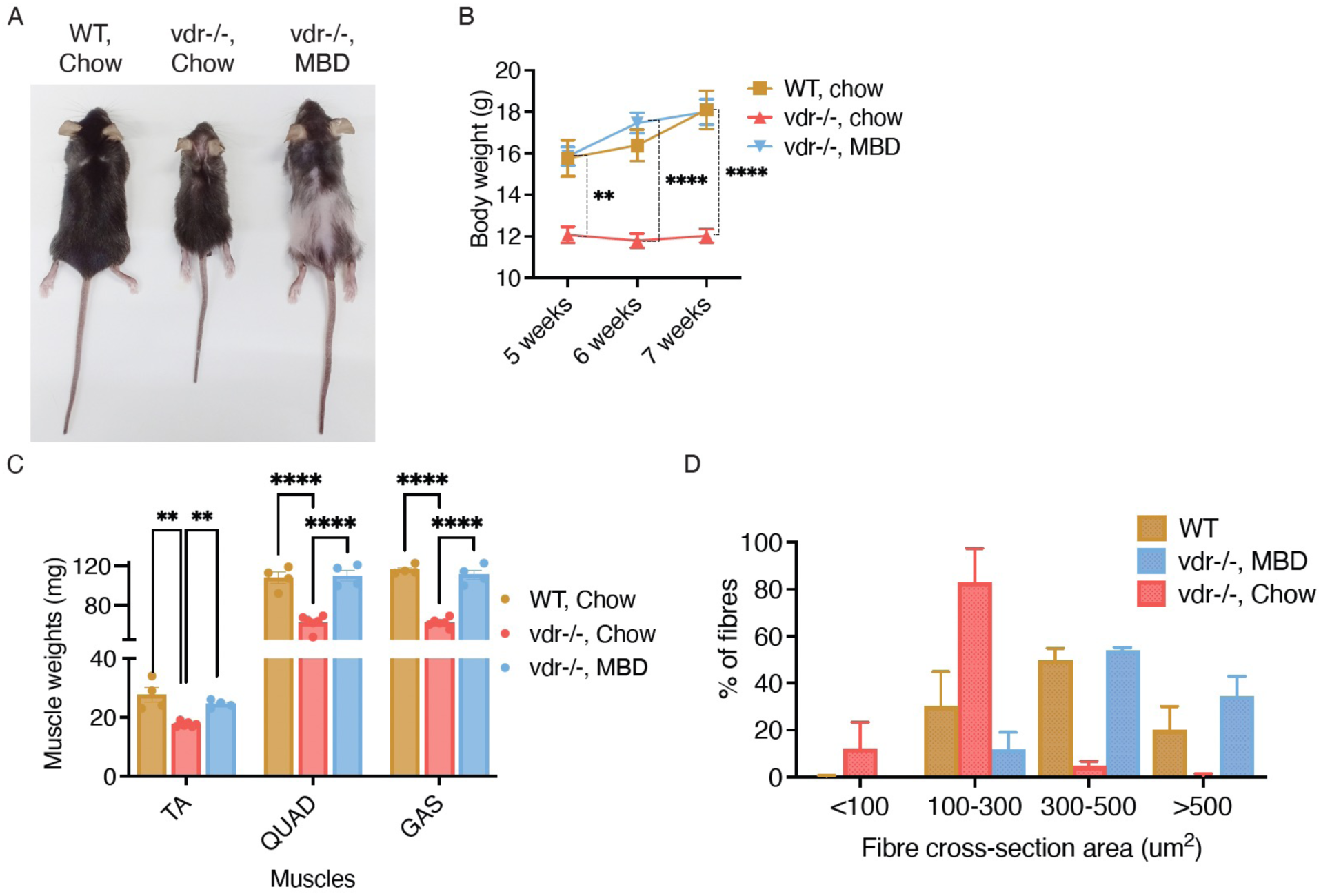
Milk based diets restore skeletal muscle atrophy in vdr-/- mice. **A.** Representative image displaying body size of WT and vdr-/- mice fed on chow, and MBD diets. **B.** Body weight (in grams) of WT and vdr−/− mice at 5, 6 and 7 weeks of age on Chow and MBD diets (*n* = 6). **C.** Tibialis anterior (TA), gastrocnemius (GAS) and quadriceps (QUAD) muscle weight of 7- week-old WT and vdr−/− on Chow and MBD diets. **D.** Quantification of muscle fiber area (gastrocnemius) of WT and vdr−/− mice at 7 weeks of age on Chow and MBD diet (n = 5). All graphs show mean ± SEM. *p < 0.05, **p < 0.01, ***p < 0.001, ****p < 0.0001 by two-way ANOVA (B, and C).

**Supplementary figure 3:**
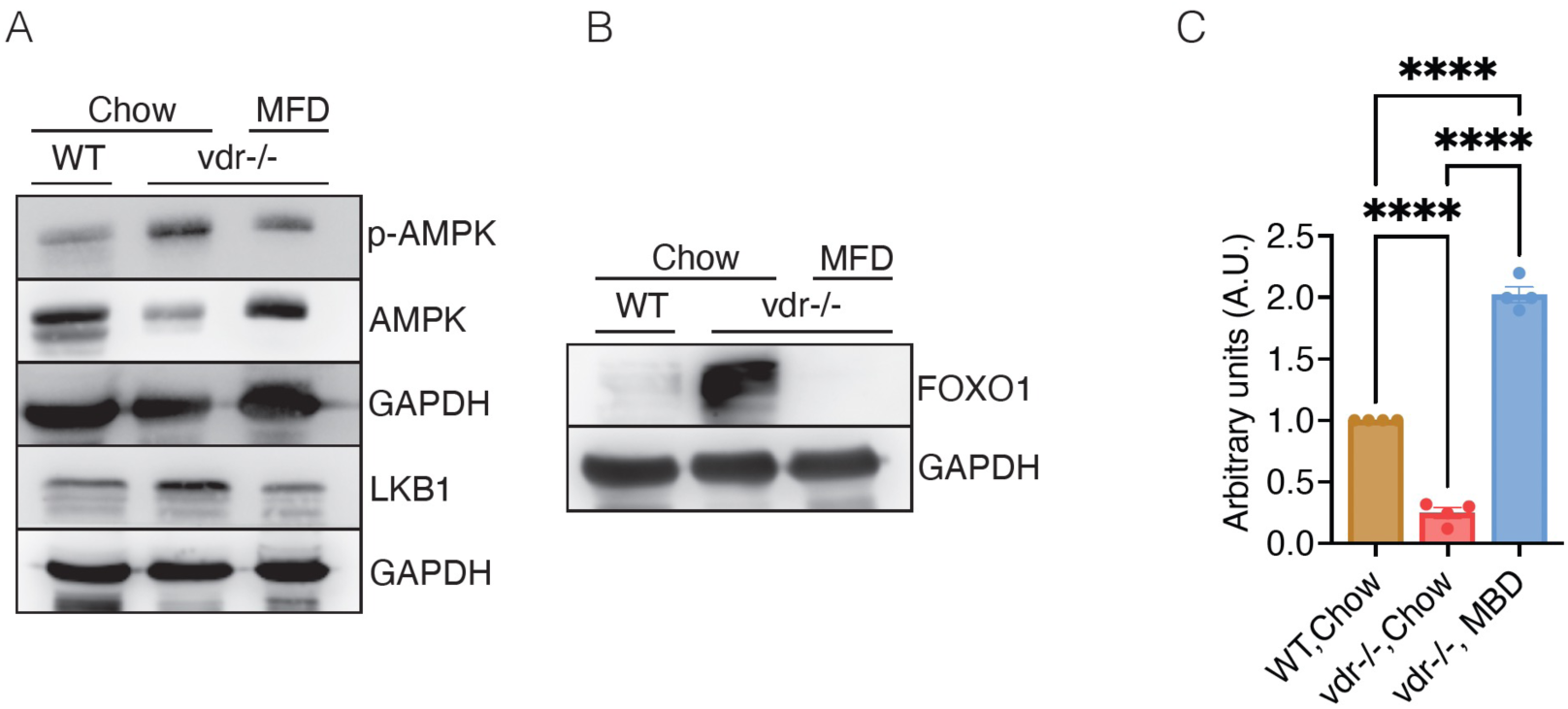
Milk-fat enriched diet restores the protein and energy homeostasis in skeletal muscles. **A-B.** Representative western blot for p-AMPK, AMPK, and LKB1 (A), and FOXO1 (B) protein levels in hindlimb of skeletal muscles of 7-week-old WT and vdr-/- mice on chow and MFD diets (n=4). **C.** Quantification of the puromycin blot represented in figure 3G. Graph in C show mean ± SEM. Statistical significance was determined by one-way-ANOVA (*: *P* < 0.05, **: *P* < 0.01, ***: *P* < 0.001, ****: *P* < 0.0001).

**Supplementary figure 4:**
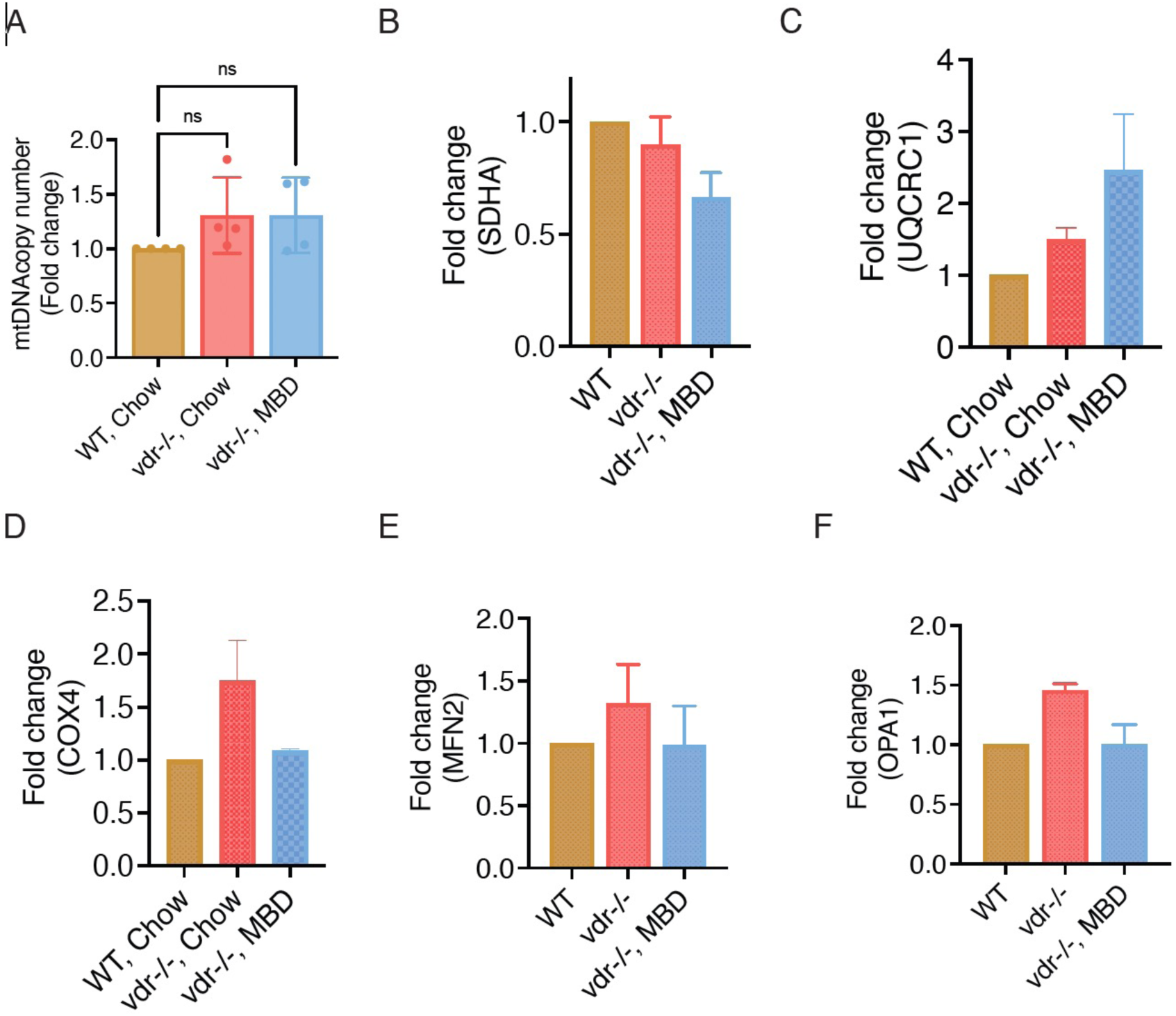
Figure 4: The milk-based diet increases mitochondrial activity and lipid metabolism in the skeletal muscles of vdr-/- mice. **A.** Fold change in mitochondrial DNA content in skeletal muscle of vdr-/- and WT mice fed on chow, and MBD diets. **B-F.** Quantification of western blots of mitochondrial proteins SDHA (B), UQCRC1 (C), COX4 (D), MFN2 (E) and OPA1 (F) in skeletal muscle of vdr-/- and WT mice. All graphs show mean ± SEM. Statistical significance was determined by one-way-ANOVA (*: *P* < 0.05, **: *P* < 0.01, ***: *P* < 0.001, ****: *P* < 0.0001).

**Supplementary figure 5:**
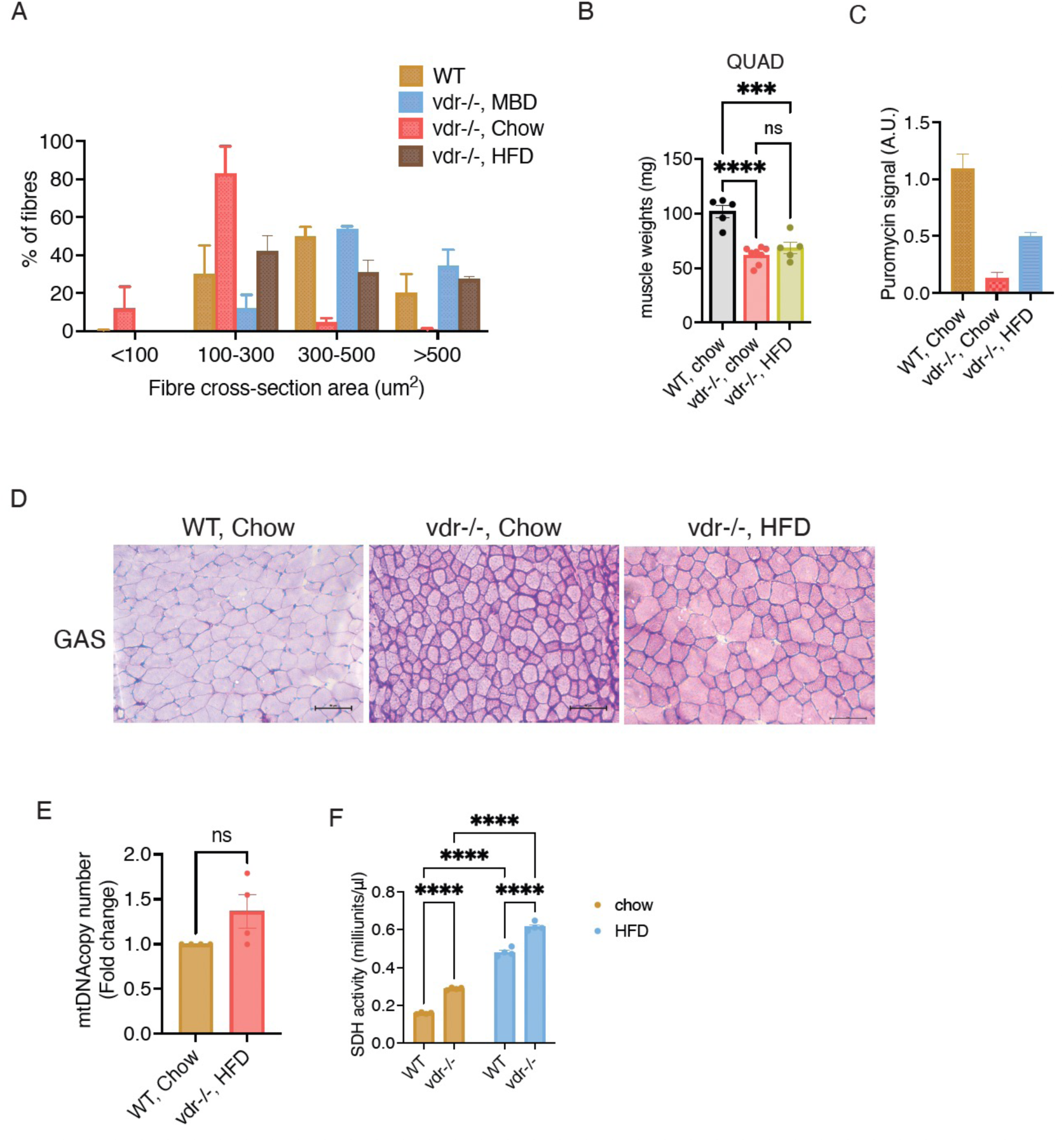
A lard-based, high-fat diet restores muscle energy levels but fails to fully restore muscle mass. **A.** Quantification of muscle fiber area (gastrocnemius) of WT and vdr−/− mice at 7 weeks of age on Chow, MBD and HFD diet (n = 5). The values used for WT, vdr-/- on chow and MBD are same as in Supplementary Figure 1 as the quantification was done with the same cohort of mice. **B.** Quadriceps (QUAD) muscle weight of 7-week-old WT and vdr−/− on Chow and HFD diets. **C.** Quantification of the puromycin blot represented in figure 5G. **D.** Micrographs of PAS stained transverse sections of GAS muscles from WT and vdr−/− mice on chow and HFD diets. 10 μm cryosections of gastrocnemius muscles were used for staining. Scale bar indicate 50μm. Magnification: 20X. **E.** Mitochondrial DNA copy number in skeletal muscle of vdr-/- and WT mice fed on chow, and HFD diets (n=4). **F.** Succinate dehydrogenase complex (complex II) activity measured in gastrocnemius muscles of WT and vdr-/- mice fed on chow and HFD diets (n=4). All graphs show mean ± SEM. Statistical significance was determined by one-way-ANOVA (B & C) or two-way-ANOVA (F) or unpaired t-test (E) (*: *P* < 0.05, **: *P* < 0.01, ***: *P* < 0.001, ****: *P* < 0.0001).

**Supplementary figure 6:**
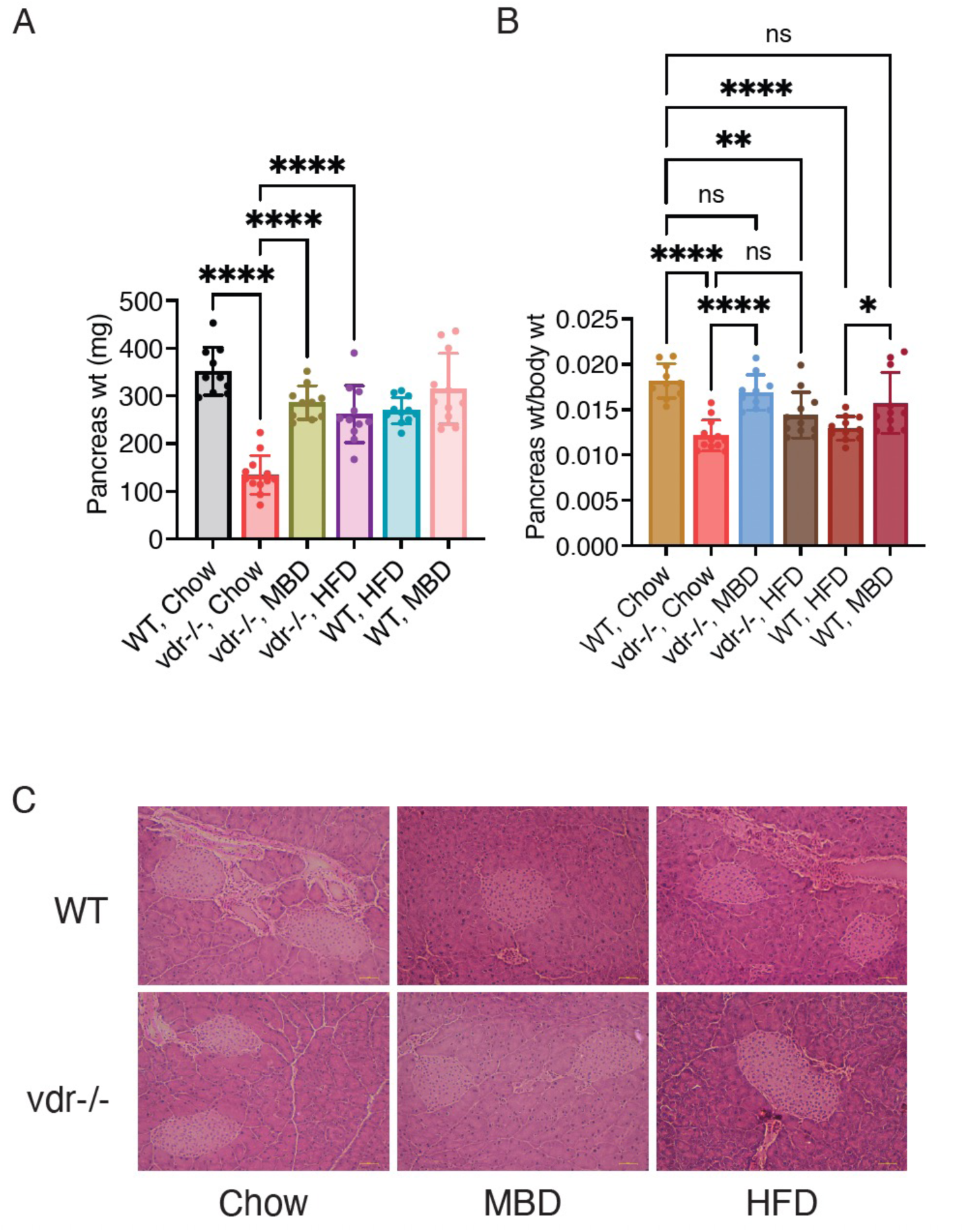
Milk-based diet could restore insulin response in vdr-/- but HFD led to inhibition of insulin synthesis. **A & B.** Weight of pancreas (A) and ratio of pancreas weight to body weight (B) at 7 weeks of age in vdr-/- and WT mice fed on chow, MBD and HFD diets. **C.** Micrographs of H&E stained paraffin embedded pancreas sections at 7 weeks of age in vdr-/- and WT mice fed on chow, MBD and HFD diets. All graphs show mean ± SEM. Statistical significance was determined by one-way-ANOVA (A & B) (*: *P* < 0.05, **: *P* < 0.01, ***: *P* < 0.001, ****: *P* < 0.0001).

**Supplementary Table 1: Comparison of different diets used in this study.** Protein, carbohydrate and fat contribution to energy for each of the diets is given.

**Supplementary Table 2: Table listing all primers used in this study.**

**Supplementary Table 3: List of antibodies used in this study.**

**Supplementary Dataset 1. Results of differential expression analysis of WT and vdr-/- QUAD muscles.** DESeq2 tool was used for DE analysis. Upregulated genes are shaded in green and downregulated genes are shaded in orange.

